# Mitigating Pandemic Risk with Influenza A Virus Field Surveillance at a Swine-Human Interface

**DOI:** 10.1101/585588

**Authors:** Benjamin L. Rambo-Martin, Matthew W. Keller, Malania M. Wilson, Jacqueline M. Nolting, Tavis K. Anderson, Amy L. Vincent, Ujwal Bagal, Yunho Jang, Elizabeth B. Neuhaus, C. Todd Davis, Andrew S. Bowman, David E. Wentworth, John R. Barnes

**Affiliations:** Battelle Memorial Institute, Atlanta, Georgia, USA; Oak Ridge Institute of Science and Education (ORISE), Oak Ridge, Tennessee, USA; The Ohio State University, Department of Veterinary Preventive Medicine, Columbus, Ohio, USA; Influenza Division, National Center for Immunization and Respiratory Diseases (NCIRD), Centers for Disease Control and Prevention (CDC), Atlanta, Georgia, USA; National Animal Disease Center, Agricultural Research Service (ARS), United States Department of Agriculture (USDA), Ames, Iowa, USA

## Abstract

Working overnight at a large swine exhibition, we identified an influenza A virus (IAV) outbreak in swine, nanopore-sequenced 13 IAV genomes from samples collected, and in real-time, determined that these viruses posed a novel risk to humans due to genetic mismatches between the viruses and current pre-pandemic candidate vaccine viruses (CVV). We developed and used a portable IAV sequencing and analysis platform called *Mia (Mobile Influenza Analysis)* to complete and characterize full-length consensus genomes approximately 18 hours after unpacking the mobile lab. Swine are important animal IAV reservoirs that have given rise to pandemic viruses via zoonotic transmission. Genomic analyses of IAV in swine are critical to understanding pandemic risk of viruses in this reservoir, and characterization of viruses circulating in exhibition swine enables rapid comparison to current seasonal influenza vaccines and CVVs. The *Mia* system rapidly identified three genetically distinct swine IAV lineages from three subtypes: A(H1N1), A(H3N2) and A(H1N2). Additional analysis of the HA protein sequences of the A(H1N2) viruses identified >30 amino acid differences between the HA1 portion of the hemagglutinin of these viruses and the most closely related pre-2009 CVV. All virus sequences were emailed to colleagues at CDC who initiated development of a synthetically derived CVV designed to provide an optimal antigenic match with the viruses detected in the exhibition. In subsequent months, this virus caused 13 infections in humans, and was the dominant variant virus in the US detected in 2018. Had this virus caused a severe outbreak or pandemic, our proactive surveillance efforts and CVV derivation would have provided an approximate 8 week time advantage for vaccine manufacturing. This is the first report of the use of field-derived nanopore sequencing data to initiate a real-time, actionable public health countermeasure.

---

Mobile sequencing technology, specifically the MinION sequencing platform from Oxford Nanopore Technologies (Oxford, UK), has successfully generated rapid genomic surveillance data on-site during outbreaks of Ebola^1^, Zika^2^, and Lassa^3^ viruses. These studies relied on transfer of raw read data to centralized institutions harboring high-performance computational resources to perform extensive phylogenetic analyses. These analyses resulted in a deep understanding of the viral evolution before and during the outbreak, as well as estimation of transmission chains. Their reliance on vast databases and computationally intensive algorithms limited their ability to influence real-time prevention strategies and potentially stop further transmission events. Furthermore, the lack of vaccines to these viruses meant that the real-time derived sequence data could not be leveraged to identify a suitable vaccine. On the other hand, influenza viruses have multiple enzootic reservoirs, cause annual epidemics in humans, and rapidly evolve, which necessitates constant genomic surveillance and iterative vaccine development. Given this need, we have an opportunity to mitigate the risk of an influenza outbreak at its source in real-time by analyzing the viral genomes present and, subsequently, developing the best matched candidate vaccine virus (CVV).

Influenza A viruses (IAV) circulating in swine have the potential to infect humans and are capable of causing pandemics^4^, as evidenced by the 2009 H1N1 pandemic (**Figure 1**). Since the 2009 H1N1 pandemic, there have been 464 human cases of swine-origin influenza viruses (termed variant influenza viruses) resulting from exposure to swine, most commonly during attendance at agricultural fairs. The worst outbreak since 2009 occurred during the summer of 2012 and resulted in at least 306 H3N2 variant cases across 10 US states^5^. These cases were primarily pediatric, with half of the cases under the age of 7. While the majority of post-2009 pandemic variant virus infections resulted in mild influenza-like illness (ILI), 16 pediatric hospitalizations and one pediatric fatality were reported. Since the majority of variant virus infections have been detected in children less than 18 years of age^6^, there is an increased risk for onward transmission in day-care settings and schools if a variant virus possessed the ability for human-to-human transmission. Thus far, human-to-human transmission of known variant outbreaks has been limited based on available epidemiologic data. However, due to the sporadic nature of these infections, the immunologically naïve population affected, and the persistent circulation of evolutionarily diverse IAV in the swine host, the Centers for Disease Control and Prevention and other World Health Organization (WHO) Collaborating Centers for Influenza and Essential Regulatory Laboratories have developed pre-pandemic candidate vaccine viruses (CVVs) that target specific swine influenza virus subtypes and HA gene lineages that have caused variant virus infections^7^. To date, eleven CVVs representing antigenically diverse groups of swine influenza viruses have been proposed or already developed for pre-pandemic preparedness purposes^8^. Ongoing surveillance efforts to assess the suitability of existing CVVs focus on the genetic and antigenic characterization of variant viruses detected in humans, but may also rely on characterization of viruses identified in swine in order to forecast which viruses are more likely to circulate at the animal-human interface and result in future human infections. This work involved sampling a large exhibition swine population in order to detect and characterize swine viruses in support of surveillance and forecasting activities.

**Figure 1.**
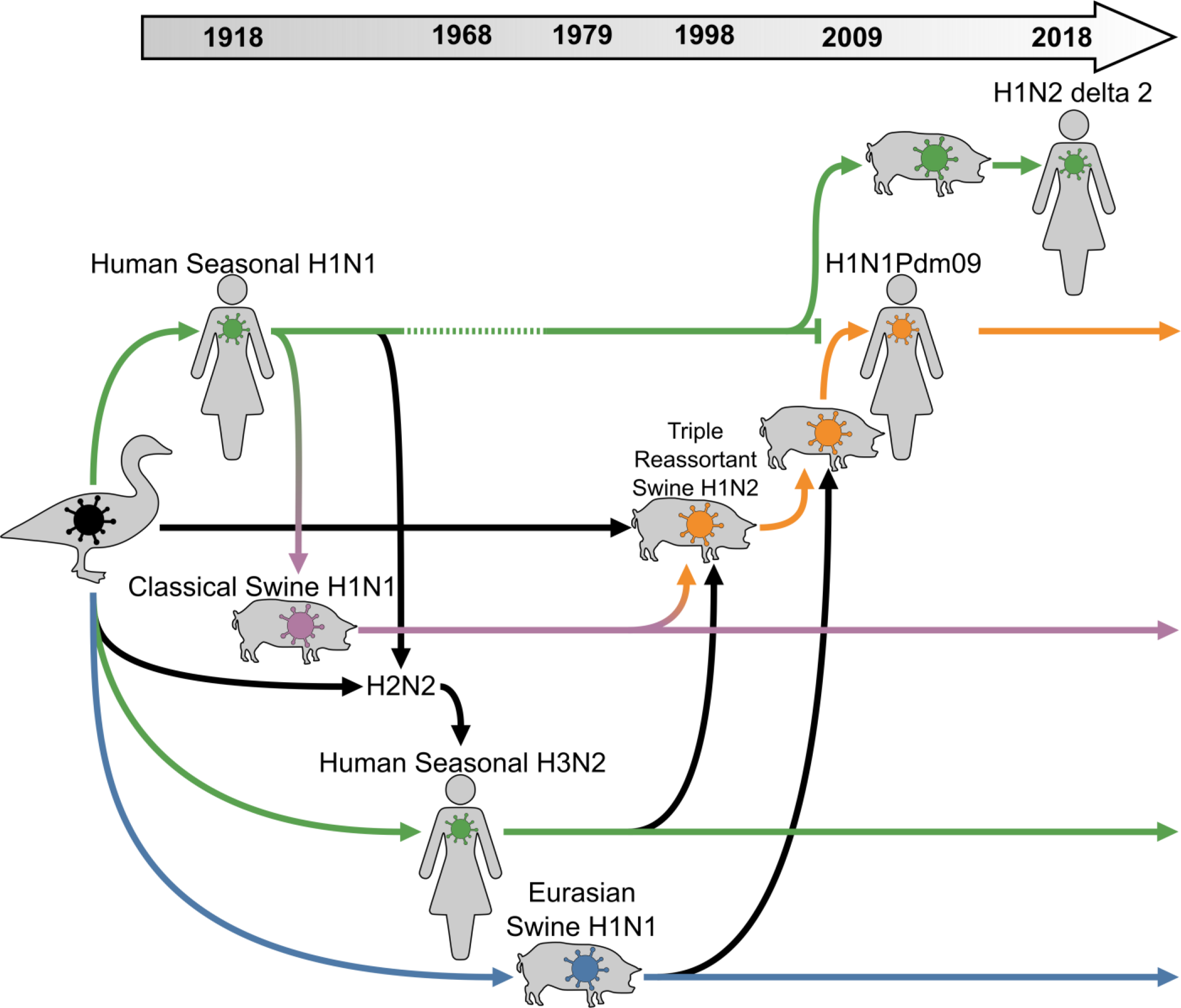
Genetic relationship of human and swine virus lineages. Colored arrows follow the evolution of hemagglutinin. Human seasonal lineages (HSL) are green, classical swine North American lineage H1N1Pdm09 (CSNAL H1N1Pdm09) is orange, classical swine North American lineage (CSNAL) is purple, and Eurasian swine lineage (ESL) is blue. In 2009, influenza A/H1N1Pdm09 virus displaced the previous human seasonal A/H1N1 virus from humans which continues circulating in swine. This virus reassorted, picking up the N2 segment from the Triple Reassortment viruses into the A/H1N2 δ2 lineage that caused human infections in the summer of 2018. The history of this HA can be traced back to the avian virus that caused the 1918 pandemic, through extinction and reemergence, and through decades of human seasonal activity.

Exhibition swine have a greater risk for viral spillover into humans as compared to production swine due to their close and protracted interaction with people and their nation-wide travel^9^. Exhibitors travel with their swine across geographical regions in the US to attend national, state, and local exhibitions. At these exhibitions, swine of various ages and from many geographical regions intermingle. Throughout the exhibition, swine are walked around the barns, shuttled into staging pens, and exhibited in a corral simultaneously with a dozen or more swine. The swine are typically shepherded by children as many exhibition rules require that individuals presenting a swine within the exhibition ring be under the age of 20. The exhibition itself can last up to a week, during which time swine are living in immediate proximity to others and have direct contact with humans. Children, in particular, may be immunologically naïve to swine influenza viruses that are genetically and antigenically distinct from recent seasonal influenza viruses or vaccines to which they have been exposed^6^. The increased risk at this unique interface warrants concerted proactive surveillance and simultaneously serves as a proving ground for new real-time in-field genomic approaches to help prevent zoonotic infections and potential outbreaks. Due to the lengthy process of collecting samples in the field and shipping them to centralized laboratories, genome sequencing and analysis often occurs several weeks or months after sample collection. Thus, the current surveillance process is less able to contribute real-time data during the emergence of pandemic threats^10^. Here, we describe the development and deployment of a rapid and portable IAV sequencing pipeline we call *Mia* [miə, MEE-uh]: *Mobile Influenza Analysis* and genomic results obtained from this surveillance.

## Results

### Development of the *Mia* Platform

We created a mobile influenza genomics platform to estimate the risk to humans posed by viruses within a local IAV outbreak by performing in-field extraction and amplification of influenza A viruses, complete genome sequencing and automated genome assembly, BLAST comparisons, phylogenetic analysis and variant detection to candidate vaccine viruses. We used our high-throughput influenza virus sequencing pipeline^11^ as a template for developing *Mia* (**Figure 2**). Our goal was a rapid workflow in a compact platform that is easily transported by two people on a commercial airliner. For RNA extraction, we elected to use Akonni Biosystems’ TruTip Rapid RNA kit due to its small footprint and demonstrated effectiveness on influenza A viruses ^12^. We chose the mini8 by MiniPCR due to its small size, ability to run 8 samples at a time, low cost, low power needs, and overall simplicity. Due to the high fidelity of our rapid amplification strategy ^11^, we were able to simplify amplicon quality control from fragment analyzation to fluorometric quantification, which is already a necessary step for amplicon pooling. For sequencing, we chose the nanopore sequencing platform MinION for the extremely small size, rapid sequencing speed, and live access to data as it is sequencing. The final inventory is available in **Table S1**.

**Figure 2.**
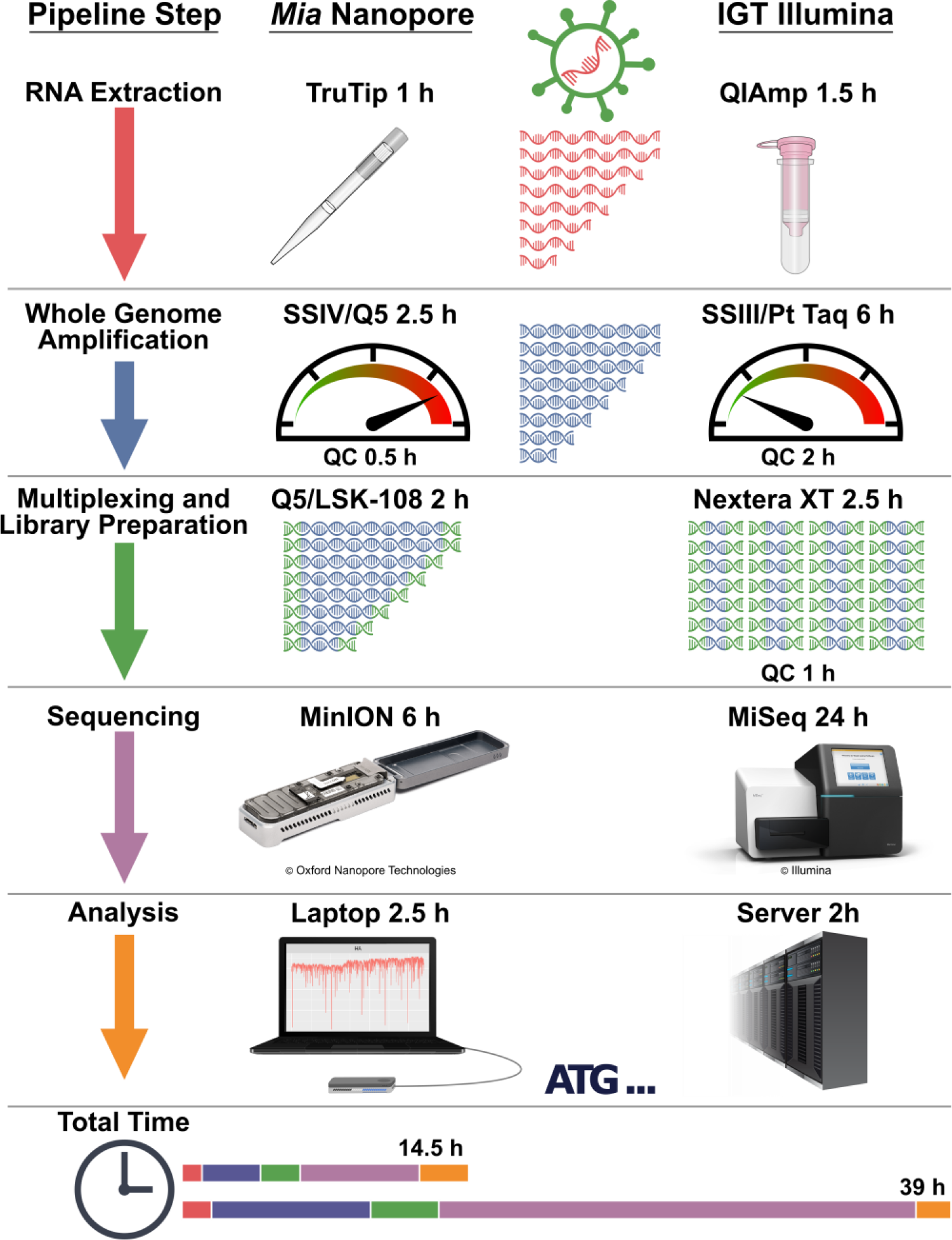
Timeline comparison of portable versus centralized surveillance pipelines. The *Mia* pipeline is modeled after the IGT pipeline with special considerations for speed and portability and is expected to get results from 24 samples in a maximum of 14 hours and 30 minutes. The IGT pipeline is designed for throughput and accuracy and would get results from 96 samples in 39 hours if operated continuously.

### Sample Processing

Our surveillance target was a large agricultural event featuring exhibition swine. The swine began arriving on Sunday (day 0) and received an initial veterinary screening upon entry. We began scouting swine on day 3 for influenza-like illness (ILI) and noted the locations in the exhibition barn of animals with clinical signs of respiratory disease. During the day, the barns were very crowded with swine, presenters, their families, farm hands, visitors and event staff. To avoid interfering with the event’s proceedings, the swabbing and sequencing was conducted during the event’s off-hours. Beginning at 4 AM on day 4, we swabbed ILI-identified swine and their pen neighbors (n=94) for influenza A virus (IAV) with Flu DETECT^®^ Swine Kits. We detected seven IAV positive samples, but suspected additional samples collected from pen neighbors might be positive by sequence analysis. At 11 PM on day 4, we erected the *Mia* platform inside the exhibition barn and began the workflow on 24 samples collected from the rapid test positive swine and their immediate neighbors. We began RNA extraction at 11:40 PM, amplification of the full genome at 12:47 AM, and quantified barcoded amplicons at 4:30 AM. Sample concentrations ranged from to 12.7 to 41.9 ng/μL (**Table S2**). We normalized and pooled the barcoded samples to 1 μg total, and began library construction at 4:50 AM. The final library amounted to 279 ng.

The library was ready for sequencing at 6 AM, however the MinKNOW controller software on an Ubuntu 16.04 operating system failed to recognize multiple MinION sequencers. We were able to use a cellphone as a mobile hotspot to download and install the latest version of MinKNOW. This fix required MinKNOW and sequencing on the Windows 10 partition, which negated use of our real-time Linux-based analytical pipeline. At this time people were arriving at the barn to begin exhibition activities and to avoid disrupting the event, once the sequencing was initiated, we transported the sequencer attached to the laptop powered by the laptop’s battery to our hotel room via automobile. Following 6 hours of sequencing, we stopped the run and transferred the raw fast5 data onto the Ubuntu partition, which initiated the analyses.

### Base-calling and Genome Assembly

Sequencing produced 99,988 reads that were base-called and demultiplexed with Albacore v.2.2.7. A total of 67,105 reads passed a basic QC threshold Q-value ≥ 7 and 22,552 of these reads were assigned to one of 24 barcodes (**Figure S1**). The distribution of read lengths followed an expected pattern with peaks corresponding to IAV segment lengths (**Figure S1**). We assembled demultiplexed reads with IRMA^13^ and obtained full IAV genomes for 13 of the 24 samples, including all samples that had Ct values < 30 (Ct 30 ≃ 100 EID50, data not shown) (**Table 1**, **Table S3**, and **Figures S2-21**).

**Table 1.**
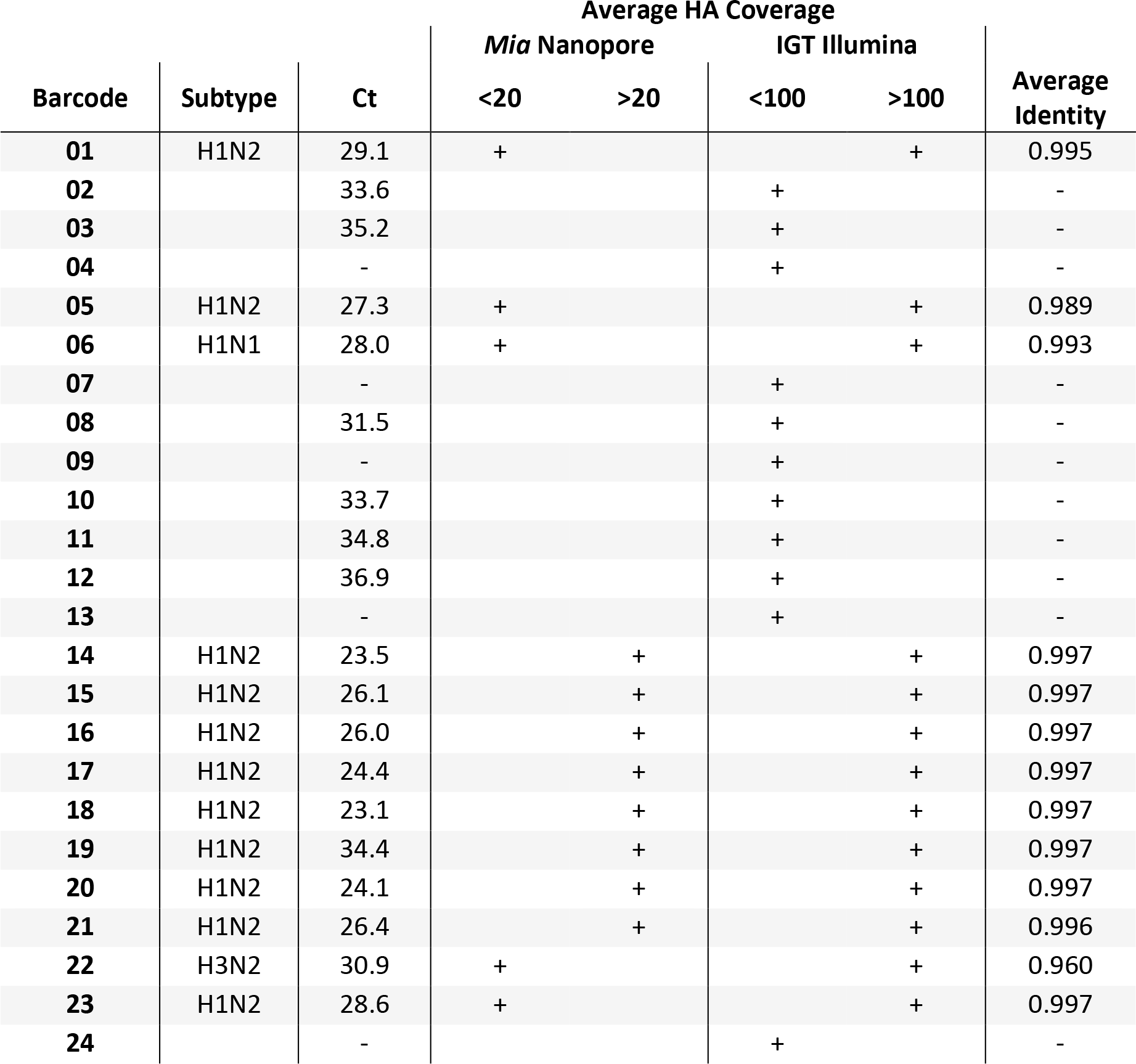
Sequencing results. HA coverage is indicated for each technique by its threshold. Samples with zero HA coverage are left blank. Complete coverage data can be found in **Table S3**. Average identity is displayed for all segments.

### Real-Time Phylogenetic Analysis

We produced automated per-segment nucleotide alignments from all sequenced samples and a reference panel of variant IAVs that were transmitted from swine to humans between 2003 and 2016. The reference panel of viruses has broad diversity including swine-origin viruses of classical swine lineages [1A.1 (H1-α), 1A.2 (H1-β), 1A.3.3.3 (H1-γ)], human seasonal lineages (1B.2.1 (H1-δ2) and 1B.2.2 (H1-δ1)), Eurasian avian-like lineages (1C.1 and 1C.2), avian-origin North American lineage, and human seasonal lineages (H3, H1 pdm09, N1 and N2)^14–17^. Each segment-alignment was automatically built into separate phylogenetic trees alongside a heat map displaying the top BLAST result per segment against the annotated references (**Figures S22-29**). Viruses collected during our investigation included 1 H3N2 (barcode 22), 1 H1N1 (barcode 06) and 11 H1N2 (barcodes 01, 05, 14-21, and 23). These 11 H1N2 viruses contained identical genome constellations. Both the HA and NA gene segments were closely related to 2017/2018 H1N2 influenza viruses circulating in the U.S. swine population and have been sporadically detected in previous A(H1N2)v zoonotic infections. The HA gene segments belonged to the delta 2 sublineage or clade 1B.2 according to the swine influenza virus H1 HA global nomenclature system. This lineage is common to swine in the U.S. and is derived from human, seasonal H1N1 virus that was transmitted from humans to pigs in the early 2000s^18^. Each of the H1N2 viruses possessed an NA gene derived from the 1998 human, seasonal H3N2 lineage now found in U.S. swine influenza viruses. This NA is a distinct gene lineage from the NA gene found in the closest CVV, A/Ohio/35/2017 (2002 human, seasonal NA lineage). The PB2, PB1, PA, and NS gene segments were of triple reassortant internal gene cassette (TRIG) origin, commonly found in swine influenza viruses in the U.S. The NP and M gene segments belonged to the 2009 H1N1 pandemic virus lineage (**Figure 3A**).

**Figure 3.**
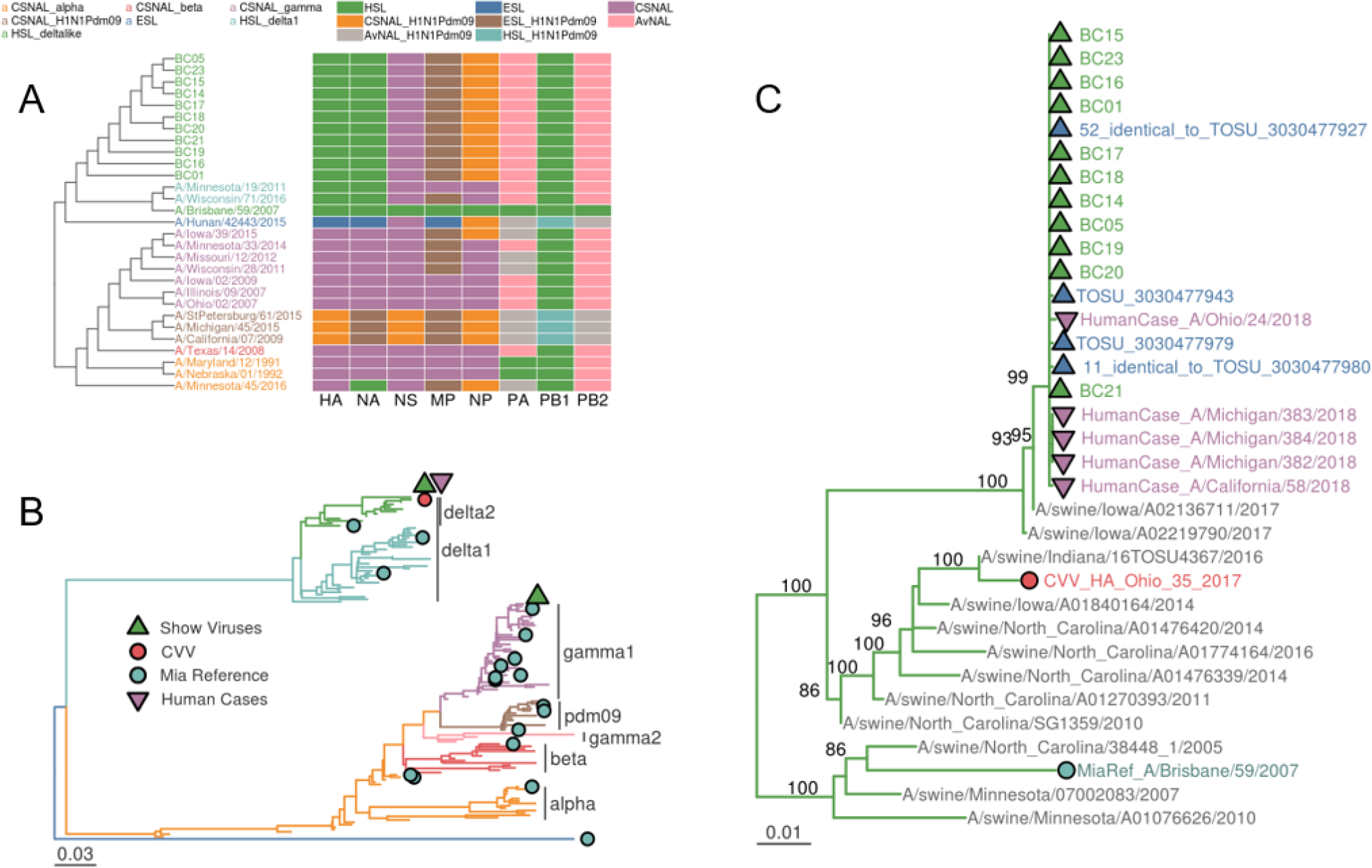
Phylogenetic relationships of HA-H1 sequences from exhibition-sampled swine, broad swine, swine-variant and late summer 2018 human cases of swine-variant IAVs. A) Recreation of the phylogenetic cladogram and genome constellation heatmap displayed in *Mia*’s Rshiny application. This automatically generated figure showed us that our sampled viruses had HA sequences most similar to the H1-delta2 A/Brisbane/59/2007 sequence and that they all shared a similar genome constellation. CSNAL = Classical Swine North American Lineage; HSL = Human Seasonal Lineage; ESL = Eurasian Swine Lineage; AvNAL = Avian North American Lineage; H1N1Pdm09 = H1N1 2009 pandemic lineage. B) Maximum likelihood phylogeny of swine IAV HA-H1 sequences incorporating the swine-variant viruses *Mia* used as references and the nearest candidate vaccine virus (Ohio/35/2017) to our found outbreak of delta-2 IAV. C) Detail of the maximum likelihood phylogeny of delta-2 HA sequences containing our field-processed IAVs (BCxx; green), 65 additional field-samples that were laboratory-processed (blue), the nearest CVV (red), *Mia* reference (light blue) and five human cases of swine-variant (purple) IAVs. Bootstrap values are annotated on internal nodes.

### Real-Time Comparison of HA1 proteins to Candidate Vaccine Viruses

Our automated real-time pipeline also provided amino acid difference tables of the mature HA1 protein sequence versus a set of WHO candidate vaccine viruses (CVV). A/Ohio/9/2015 and A/Ohio/13/2017 were the nearest CVVs to the H1N1 and H3N2 viruses and had 2 and 0 antigenic changes, respectively. These two viruses also had relatively low sequence coverage of the HA segment (**Figures S4 & S19**). A/Ohio/35/2017 was the most similar CVV in our database to the 11 H1N2 viruses and had 32 amino acid differences compared to their consensus sequence (**Figure 4**, **Figure S30**, and **Table S4)**. Seven differences were identified in predicted H1 antigenic sites, including 4 in antigenic sites A or B^19^. More generally, the H1N2 virus HA genes were only ~93% identical to the nucleotide sequence of Ohio/35/2017 and, as discussed above, had NA genes belonging to different ancestral lineages. Based on the genetic differences and markers of antigenic drift detected by the analysis, the HA and NA consensus sequences, including UTRs, were electronically transmitted to CDC’s Vaccine Preparedness Team in the Influenza Division to initiate the development of a synthetic candidate vaccine virus. We were able to start this intervention from the field roughly 18 hours after the initial setup of *Mia* (**Table S5**).

**Figure 4.**
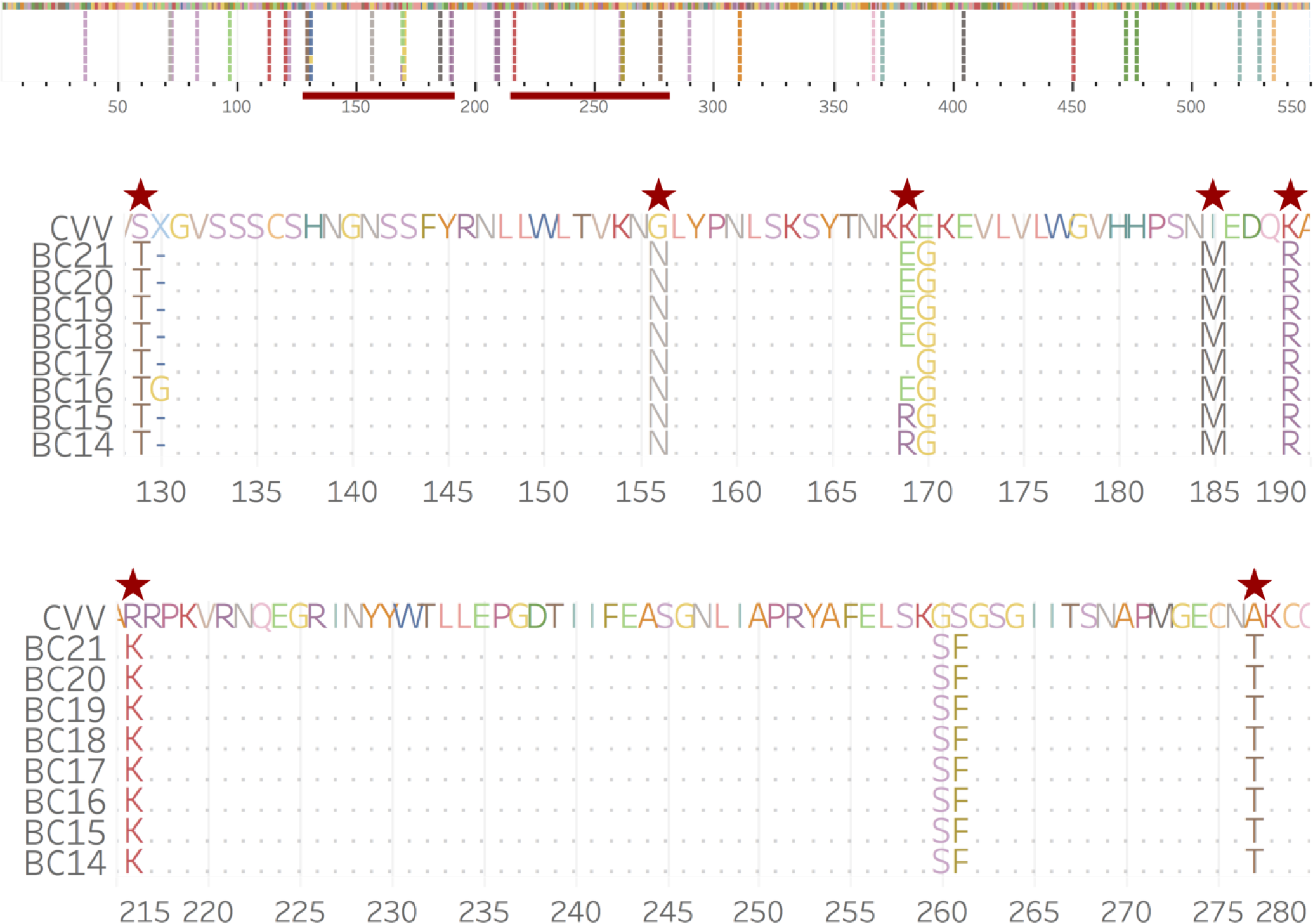
Mature H1 amino acid sequence alignment of H1 delta-2 sequences from *Mia* nanopore sequencing versus A/Ohio/35/2017, the nearest CVV. Red stars indicate antigenic sites. This figure was created post-field for publication purposes, however the same information was viewable in an alignment viewer in the field as well as in *Mia*’s dashboard table.

### Post-Field Analyses

To assess the MinION-derived sequence accuracy we transported RNA and DNA amplicons created onsite to the CDC in Atlanta, GA for confirmatory sequencing via Illumina MiSeq using the standard Influenza Genomic Team’s (IGT) surveillance pipeline^13^. On the MiSeq, 13 samples had completed genomes with average coverage ≥ 100x for each segment (IGT’s standard QC threshold). Of those 13, 12 had MinION-derived mean coverage of ≥ 5x across segments with 8 having ≥ 20x coverage across HA, the primary target of protective immune responses and the most important protein for vaccine development (**Table S3**). There was high concordance (R^2^ = 0.87; **Figure S31**) of mean segment coverage produced by MinION and MiSeq for field-derived amplicons. Consensus identities averaged 99.3% across the 13 genomes (**Table S3**).

As our real-time *Mia* database contained human-cases of swine IAV, we performed additional post-field phylogenetic analyses using the outbreak data, the *Mia* database, and additional swine IAV sequences that represent the genetic diversity and evolutionary relationships of contemporary swine IAV^14,15,20^. When compared to 126 swine H1 sequences, our *Mia* identified swine-H1N2 viruses were in a monophyletic clade with contemporary 1B.2.1 (H1-δ2) HA genes from the human seasonal lineage (**Figure 3**). These viruses predominantly pair with N2 sequences that have been circulating in swine since at least 1998 as part of a triple reassortment (TRIG) group of viruses^15,16^ (**Figures 1** and **S36**). Across all segments, MiSeq-derived data mirrors the tree topology created in real-time in the field from MinION-derived sequences (**Figures 3**, **S22-29,** and **S33-41**).

To compare the viral diversity we found in our MinION-derived viral sequences to what fully existed at the exhibition, we performed our standard IAV surveillance procedure in which we sampled 425 additional swine regardless of ILI. From these samples, we detected 136 (32%) IAV-positive specimens by RT-PCR and successfully sequenced 65 full genomes on a MiSeq. We found that the diversity of viruses collected and sequenced by more traditional surveillance methods was also detected in similar proportions by our *Mia* field-sequencing (**Figures 3** and **S33-41**).

## Materials and Methods

### Logistics

Our instrument inventory was finalized after three full practice runs and consisted of three main pieces of luggage to transport *Mia* (**Table S1**, **Figure S43**). We used a Pelican (Torrance, CA, USA) Air 1615 (internal dimensions: 75.16 × 39.37 × 23.82 cm) to transport plastic consumables, ambient temperature liquids, PPE, racks, pipettes, and power strips. This is a larger suitcase that was checked during air travel. We used a smaller Pelican 1510 (internal dimensions: 50.3 × 27.94 × 19.3 cm) to transport the electronic equipment including the MinIONs, thermocyclers, vortex, microfuge, and qubit. While this case is extremely rugged and would likely fare well against the checked baggage handling, we were concerned about damage during luggage inspection and opted to carry this onto the plane. We also carried on the laptop inside of a backpack. Our cold-chain was maintained during transportation in an Ozark Trail (Potosi, MO, USA) 26 quart certified bear-resistant cooler (internal dimensions: 27.3 × 40.64 × 22.86 cm). To simultaneously transport materials at −20°C and +4°C, the cooler was partitioned with a 7.6 cm piece of Styrofoam cut to a snug fit. All enzymes and reagents requiring storage at −20°C were stored in a cold block and surrounded with frozen cold packs in the bottom of the cooler. Primers and flowcells were stored at +4°C on top of the Styrofoam and were surrounded by refrigerated cold packs. We maintained the cold chain on site by storing reagents in a hotel refrigerator/freezer before repacking the cooler to transport the reagents to the exhibition barn. The SuperScript IV was shipped to the barn on dry ice at the insistence of the manufacturer. Since ethanol over 70% cannot be transported on a commercial airliner, it was shipped via ground transportation. For a work bench, we used a small folding table found at the event and purchased plastic table cloths, Lysol, and extra garbage bags locally. To avoid disrupting the event, we setup *Mia* inside a horse stall near the pigs and worked overnight (**Figure S44**).

### Diagnostic screening and sampling

To confirm the presence of IAV and to identify the location of outbreak clusters, we screened 94 swine displaying ILI with Flu DETECT^®^ (Bridgewater, NJ, USA) Influenza A Virus Swine rapid tests via nasal swabs. We performed the screening early in the morning to locate IAV positive swine and returned to the same location for sampling and sequencing later that evening. It should be noted that due to logistical challenges, we used standard cotton swabs instead of the provided swabs, which likely contributed to the high false negative rate of the tests^21^.

We selected 24 swine for respiratory-sample collection and sequencing which included the seven positives from the screen and their immediate neighbors. We sampled these swine via sterile gauze nasal wipes and submersion in 5 ml brain heart infusion (BHI) media^22,23^.

### RNA Extraction

Immediately after sampling, we extracted RNA from the BHI samples using the Akonni TruTip^®^ (Frederick, MD, USA) Rapid RNA kit according to the manufacturer’s instructions with minor modifications ^12^. We extracted the samples in a 96 deep well block using a manual 12 channel 1000 μL pipette. Before lysis, we diluted the 70 μL samples with 180 μL of water and spiked them with 0.5 μg of Qiagen (Venlo, Netherlands) carrier RNA. We eluted the samples in 70 μL of water that had been warmed to 75°C.

### Amplification, Barcoding, Library Preparation, and Sequencing

We amplified the full influenza A virus genome using a recently described fast multi-segment reverse transcription polymerase chain reaction (fMRT-PCR)^11^ with some modifications for portability. We used three miniPCR™ (Cambridge, MA, USA) mini8 thermocyclers under laptop control for elution buffer heating, fMRT-PCR, barcoding, and library preparation. Following barcoding, we quantified the products using a Qubit™ (Thermo Fisher Scientific, Waltham, MA, USA) DNA BR/HS assay. The quantification data allowed us to normalize and pool the barcodes and served as our only quality control check during the molecular workflow. Once pooled, we prepared the amplicons for nanopore sequencing using the SQK-LSK 108 library kit (Oxford Nanopore Technologies, Oxford, UK). This protocol required cooling to 20°C; however, the temperature in the barn was roughly 30°C, and the fan-cooled mini8 thermocyclers could not get below ambient temperature. To solve this issue, we placed the thermocycler on a bed of ice, which effectively lowered the ambient temperature and allowed the mini8 to cool to 20°C. We sequenced the pooled samples on a MinION Mk 1B nanopore sequencer using flowcell FAH58363 equipped with R9.5.1 chemistry.

### Real-Time Analyses

For real-time analyses, we used a high performance Dell (Round Rock, TX, USA) Precision 7720 laptop with 64GB ram, four-core Intel Xeon 3.00GHz CPU E3-1505M v6 (2 threads/core) and two 1TB solid state hard drives, partitioned separately into a Windows 10 OS and Ubuntu 16.04 OS. *Mia*’s custom analytical application runs on the Ubuntu partition and initiates automatically via crontab upon MinION read file (fast5) origination during active sequencing. We base called and demultiplexed fast5 read files with Albacore v.2.2.7 (Oxford Nanopore Technologies, Oxford, UK) and assembled subsequent fastq files into consensus sequences with IRMA v.0.6.7^13^. We used a custom MinION configuration module for IRMA that changes the default ‘FLU’ module parameters as follows: dropping the median read Q-score filter from 30 to 0, raising the minimum read length from 125 to 150, raising the frequency threshold for insertion and deletion refinement from 0.25 to 0.75 and 0.6 to 0.75 respectively, and lowering the Smith-Waterman mismatch penalty from 5 to 3 and the gap open penalty from 10 to 6. We checked consensus sequences against a local database containing clade-annotated IAV sequences with BLASTn v.2.7.1+^24^ and used top BLAST matches to interpret a sample’s IAV genome constellation. We aligned consensus sequences against reference sequences with MUSCLE v.3.8.31^25^. We built phylogenetic trees with FastTree v.2.1.8 Double Precision^26^ with a generalized time-reversible model, branch lengths optimized under a gamma distribution, and local support values produced from bootstrap sampling 10,000 times and visualized the trees in R v.3.4.4 with GGtree^27^. We determined amino acid differences to each candidate vaccine virus sequence in our database with a custom Python script that translates DNA consensus sequences and MUSCLE aligns the mature HA1 protein sequences. Analytical and processing status results were fed into a SQLite v.3.11.0 (Hwaci Inc. Charlotte, NC) database and visualized in a web browser with an interactive R-shiny^28^ application.

Reads are processed in 4,000-read batches for the first 5 iterations, followed by 20,000-read batches for another 5 iterations and then 40,000-read batches until reads are no longer produced by the sequencer. This stepwise batching provides the user an understanding of the run’s quality within minutes of sequencing, allowing the user to decide if it is worth continuing the run while maximizing computational resources. Results seen within the first hour of sequencing track across a 6- or even 48-hour run, with the extended run time increasing coverage and resolving a scant number of consensus bases ^11^.

### Post-Field Analyses

We transported field-derived RNA and amplicons on dry ice back to the CDC in Atlanta, GA and processed them with the Influenza Genomics Team’s standard Illumina (San Diego, CA, USA) MiSeq influenza surveillance pipeline. We processed and assembled reads with IRMA’s FLU-sensitive module ^13^ and calculated the identity of MinION-derived consensus sequences based on alignment to the sample’s MiSeq-derived consensus, considered the gold standard. Out of 425 swine sampled through our standard surveillance efforts at the exhibition, 136 tested positive for IAV by RT-PCR. Complete genome sequencing via our standard Illumina pipeline was attempted on 79 of those and produced 65 full genomes.

We performed phylogenetic analyses with MiSeq produced sequences and an annotated set of swine-IAV sequences. The annotated dataset was constructed by downloading all publicly available swine IAV data collected in North America from the Influenza Research Database^29^ on June 1, 2018. Each gene was aligned with MAFFT v7.294b^30^ and the best-known maximum likelihood phylogeny for each alignment was inferred using IQ-TREE v1.6.2^31^, with the model of molecular evolution automatically selected during the tree search. Each gene was classified to evolutionary lineage and/or phylogenetic clade^15,17,20^. Subsequently, we selected 3-10 viruses from each named clade that captured the evolutionary relationships among clades, and that represented the major HA, NA, and whole genome constellations circulating in US swine^15,20^. We then aligned the MiSeq and annotated sequences with MUSCLE v.3.8.31^25^ and inferred maximum likelihood trees using IQ-TREE^24^ v.1.6.8 implementing a generalized time-reversible model of nucleotide substitution with gamma-distributed rate heterogeneity across sites while allowing for a proportion of invariable sites and 1000 ultra-fast bootstrap approximations^32^ (iqtree −m GTR+G+I −bb 1000).

## Discussion

We successfully developed a mobile NGS suite called *Mia* and deployed it to a high priority swine-human interface that resulted in our identification of an influenza A virus outbreak. The data generated indicated that while multiple lineages were co-circulating, available candidate vaccine viruses (CVVs) were less likely to be adequate antigenic matches if the predominate viruses were to emerge in humans. Therefore, the sequence data was used to initiate synthesis of an optimal CVV within 18 hours of unpacking *Mia*. This proactive strategy of identifying the predominant swine influenza viruses in exhibition pigs prior to the start of annual agricultural fairs in the USA proved to be very useful as zoonotic infections caused by the predominate A(H1N2) virus found in exhibition pigs were detected in humans exposed at fairs in July and August 2018^33^.

We created *Mia* by selecting and optimizing laboratory equipment that fits into two standard sized suitcases that can be set up and operated in the field by two people to produce a high-quality multiplexed NGS amplicon library in 7 hours. Also in the field, we performed base calling, quality control, genome assembly, phylogenetic analysis, BLAST searching, and amino acid comparisons to current CVVs without an internet connection. We were able to monitor all these results via our custom automated application on a laptop. The numerically dominant subtype we detected were H1N2 swine influenza viruses that were genetically different from the most similar WHO pre-pandemic CVV. We determined that these viruses posed a risk of causing disease in a young population, as similar H1 viruses disappeared from seasonal circulation in humans in 2010 and were replaced by the 2009 H1N1 pandemic virus. Approximately 18 hours after sampling, we emailed consensus sequences of these viruses to CDC for immediate synthesis of a new H1N2 CVV.

By returning RNA from the field, we were able to confirm the sequencing results with our Illumina sequencing pipeline ^13^. Without considering a coverage threshold for the nanopore data, we sequenced 13 full genomes in the field. These same 13 samples were also successfully sequenced in the lab on an Illumina MiSeq. This, at least partially, confirmed the sensitivity of our field based method. By comparing MiSeq and MinION produced sequencing, we saw that the maximum MinION-derived sequence accuracy is achieved between 10-20x mean fold coverage (**Figure S32**). If we apply a 20x coverage threshold to the nanopore data, then only 4 genomes were successful in the field, but with 8 HA segment sequences passing. Importantly, the HA sequence is the most critical for making vaccine strain determinations. In future experiments, we can ensure coverage thresholds are met by monitoring coverage in real-time and more deliberately deciding when to stop sequencing, rather than simply sequencing for a defined 6 hours as we did here. Illumina sequencing also allowed us to confirm the accuracy of our in-field generated consensus sequences (99.3% overall). This level of accuracy confirms the validity of the suite of analyses that we performed in the field with the nanopore data.

While speed is a primary success of our work, there remains room for improvement. A major delay in this effort was an Oxford Nanopore Technologies’ software controller (MinKNOW) issue that failed to recognize and connect to the sequencer. The fix for this took 34 minutes and required sequencing on the Windows partition of the laptop whereas the analysis software runs on the Linux partition. This resulted in our automated analysis pipeline not having access to the raw sequencing data until we terminated the sequencing experiment after 6 hours and transferred the data across the partition. This issue has since been resolved and would now save us up to 6 hours on the entire workflow, depending on the read generation rate of a given sequencing run. We are also developing a containerized version of *Mia* that will be able to run on any operating system that supports Docker (Docker Inc, San Francisco, CA), including Windows and Mac. Moreover, we are now incorporating the use of Oxford Nanopore Technologies’ (Oxford, UK) MinIT portable GPU base caller that will dramatically reduce the time of base calling, our analytical pipeline’s tightest bottleneck.

Direct RNA sequencing of IAVs has recently been demonstrated, and while this technique is extremely fast (<2 hrs from RNA to sequencing), it is not yet suitable for clinical applications due to a lack of sensitivity and multiplexing^34^. We are exploring molecular techniques to overcome these challenges. The sequencing time required to achieve our desired 20x mean coverage across each segment is also dependent on factors which are poorly understood, though likely related to reagent stability and the integrity of individual pores on the flowcell. While this in-field trial produced nearly 100k reads in 6 hours, we have commonly produced over 1M reads in this time frame in the laboratory. Whether the low yield found in this field trial is due to the field conditions or an outlier is the subject of further investigation as we prepare for future field deployments.

The success of *Mia* hinged on our ability to curate a small reference set of virus genomes with lineage annotations per segment. While the rapid growth of viral genome sequences in public databases has allowed fine-grained phylogenetic analyses of viral pathogens, analyzing the complete evolutionary history is not necessary to identify pathogens and guide a public health intervention during an outbreak. *Mia* demonstrates that curating a small and diverse reference database is sufficient for estimating the risk of an influenza outbreak. Importantly, the reference set can be adapted for other IAV outbreak targets, such as avian IAV in live bird markets. Furthermore, *Mia*’s analytics are 100% automated as reads originate on the file system of a machine connected to the MinION. This is a critical feature in our goal of decentralizing influenza virus surveillance and outbreak response around the world, as pertinent information is provided back to the user in real-time.

Influenza A viruses (IAV) are a perpetual threat to global health security, both as human-endemic seasonal viruses and the more insidious pandemic viruses. IAV pandemics result from zoonotic transmission of IAVs to humans followed by onward human-to-human spread in populations lacking sufficient immunity. The swine-human and avian-human interfaces are of keen interest for future in-field surveillance efforts due to the pandemic risk of avian and swine influenza viruses that circulate in these settings^35,36^. *Mia* will be a useful tool to enhance the current centralized surveillance framework and can be deployed into resource poor settings and operated by two technicians. This technology can serve to bolster IAV surveillance by monitoring important transmission interfaces for emerging viruses that have pandemic potential.

## Acknowledgements

Research reported in this publication was supported by the office of Advanced Molecular Detection (AMD CAN 939018 C) at the Centers for Disease Control and Prevention. The Council of State and Territorial Epidemiologists supported this project with funds provided by the Centers for Disease Control and Prevention under Cooperative Agreement No. 5U38OT000143-05. Nasal wipes were collected as part of influenza A virus surveillance supported by the Centers of Excellence for Influenza Research and Surveillance, National Institute of Allergy and Infectious Diseases, National Institutes of Health, Department of Health and Human Services contract HHSN272201400006C. We thank bluegrass legend Mindy Barringer from CDC’s Division of Communication Services for Figure 2 visual display. Mention of trade names or commercial products in this article is solely for the purpose of providing specific information and does not imply recommendation or endorsement by the USDA. USDA is an equal opportunity provider and employer.

The authors declare no competing interests.

