## SupplementalFigures for "Mitigating Pandemic Risk with Influenza A Virus Field Surveillance at a Swine-Human Interface"

Running Title: Portable Influenza A Virus Surveillance

†B.L.R.-M. and M.W.K. contributed equally to this work

### Logistical Improvements

Logistically, there are several improvements to this strategy that can immediately be implemented for this or a more challenging scenario that may not have readily available electricity, refrigeration, or ice. At the swine exhibit, our power needs were met by a reliable 120V AC outlet; however, this would likely not be available in future efforts to deploy this technology. While all the power needs for this equipment are relatively low, this is still a major challenge and should employ a robust approach to ensure electricity is not an issue. Lithium ion batteries designed to charge cell phones have enough capacity to run our thermocyclers, microfuge, and Qubit, however charging the laptop and or providing multiple days' worth of power would require batteries too large to transport on a commercial air liner. A possible workaround for this would be to use a battery array that is not technically considered to be a single battery. Portable solar panels, while low in power, could be used to extend the life of these batteries. A second approach to electricity is to transport a small generator. Unopened generators can be checked on commercial airliners. This approach would be quite effective with the assumption that fuel can be located on site and the caveat that the generator cannot be transported back once fuel has been added. A backup strategy would be to bring a power inverter for 12V lead acid batteries.

There were two reagents that weren't transported on the commercial airliner: 95% ethanol and superscript IV. Ethanol is limited to 70% on commercial airliners and our workflow required 95% ethanol for the extraction and 80% ethanol for the bead washes. Since our samples are diluted with water before ethanol is added, we can simply combine these reagents ahead of time and add 555 µL of 65% ethanol to have the same effect. Lowering the concentration of ethanol for the bead washes would simply mean accepting a potentially lower yield and could be overcome by overloading the input material. Superscript IV is highly temperature sensitive and the manufacturer insisted it be transported on dry ice, despite recommending laboratory storage at -20°C. Dry ice can be transported on a commercial airliner, 3 or 5 lbs according to an airport ticket agent or the FAA respectively, and we can simply add a cooler to transport this reagent. Alternatively, since our cooler maintained -20°C well, we can also transport this in the cooler as designed. Continued improvements, such as these, will be implemented as *Mia* continues to be deployed to less forgiving environments.

### Supplemental Table Legends

The supplemental tables are available as a separate excel file.

**Table S1**
Complete inventory of *Mia* with storage locations

**Table S2**
Barcoded amplicon concentration for QC and pooling

**Table S3**
Sample information and sequencing results

**Table S4**
Amino acid differences between *Mia* nanopore results and the nearest CVV (A/Ohio/35/2017)

**Table S5**
Actual timeline of the *Mia* deployment from opening the cases in the barn to sending the sequences for vaccine synthesis

**Table S6**

Accessioned Data

### List of Supplemental Figures


In-Field Screenshots: S1-S30

1. Albacore Summary

Coverage by barcode

2. BC01
3. BC05
4. BC06
5. BC07
6. BC08
7. BC09
8. BC10
9. BC11
10. BC13
11. BC14
12. BC15
13. BC16
14. BC17
15. BC18
16. BC19
17. BC20
18. BC21
19. BC22
20. BC23
21. BC24

In-field trees

22. HA
23. NA
24. PB2
25. PB1
26. PA
27. NP
28. M
29. NS

30. CVV comparison

Post-field analysis

31. MinION vs MiSeq coverage
32. MinION coverage vs identity to MiSeq

Post-field phylogenetic analysis

33. HA H1 gamma
34. HA H3 2010 human like
35. NA N1
36. NA N2
37. PB2
38. PB1
39. PA
40. NP
41. M
42. NS

Photographs

43. *Mia* contents and cases
44. *Mia* mid workflow in barn

### Supplemental Figures

#### Figure S1

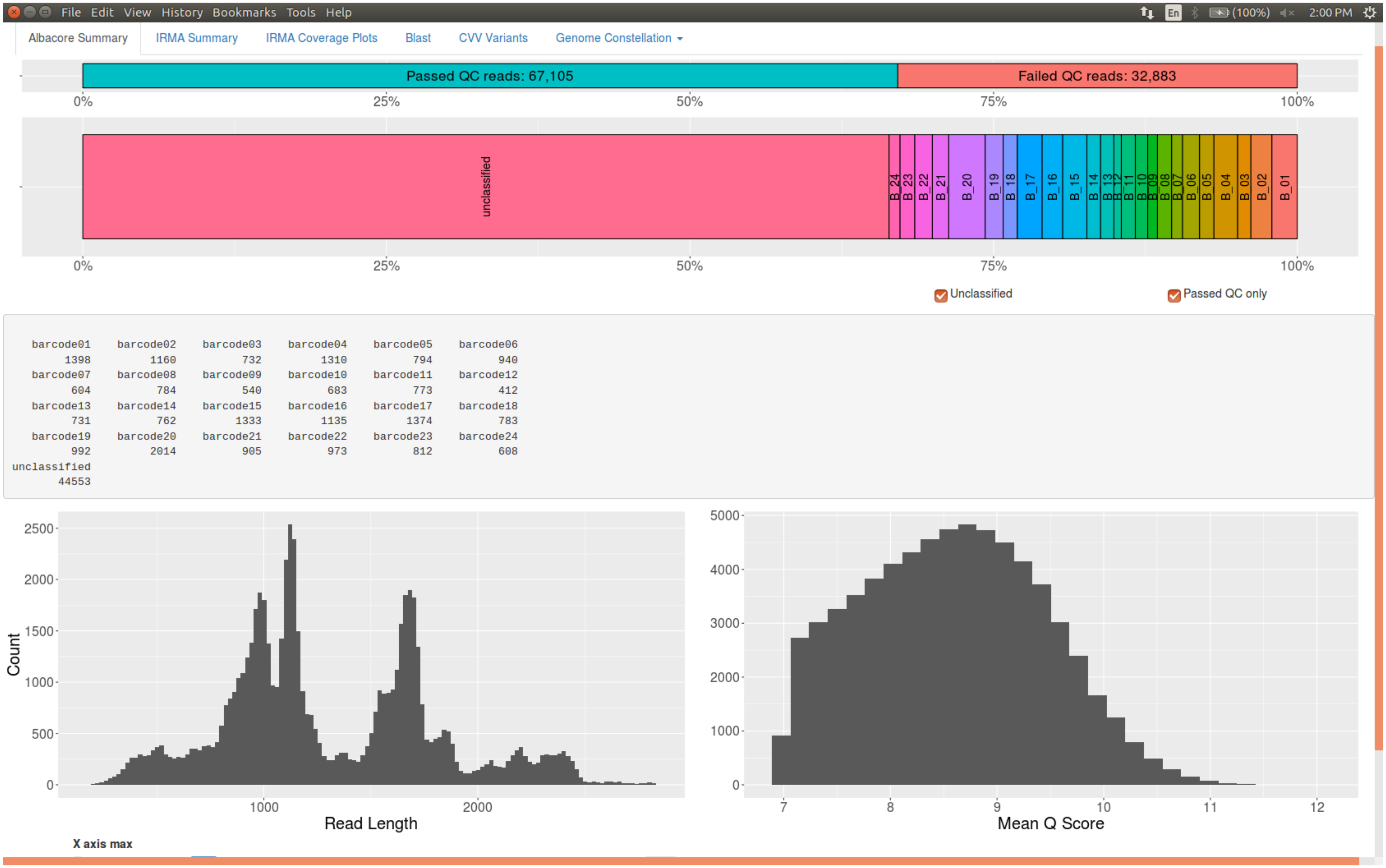


Albacore results summary screenshot

**Figure S2**

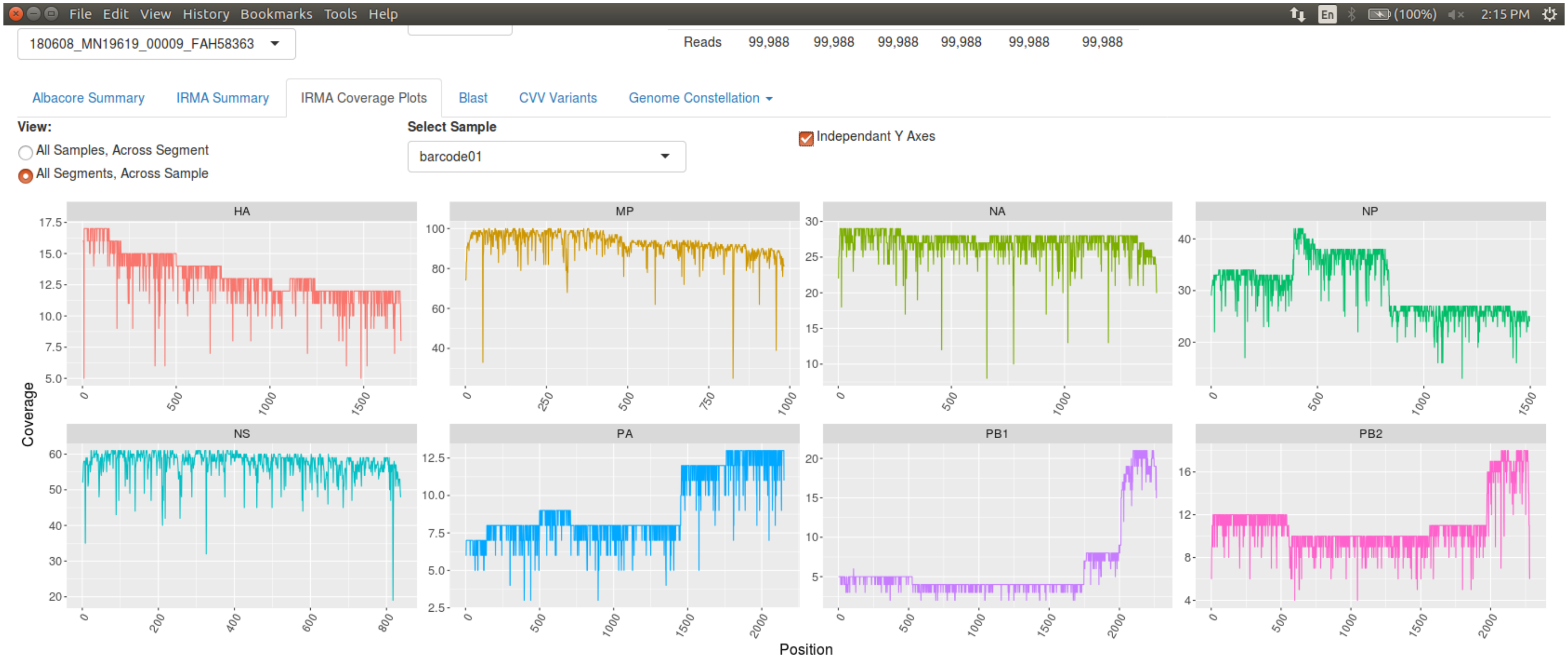


BC01 coverage screenshot

### Figure S3

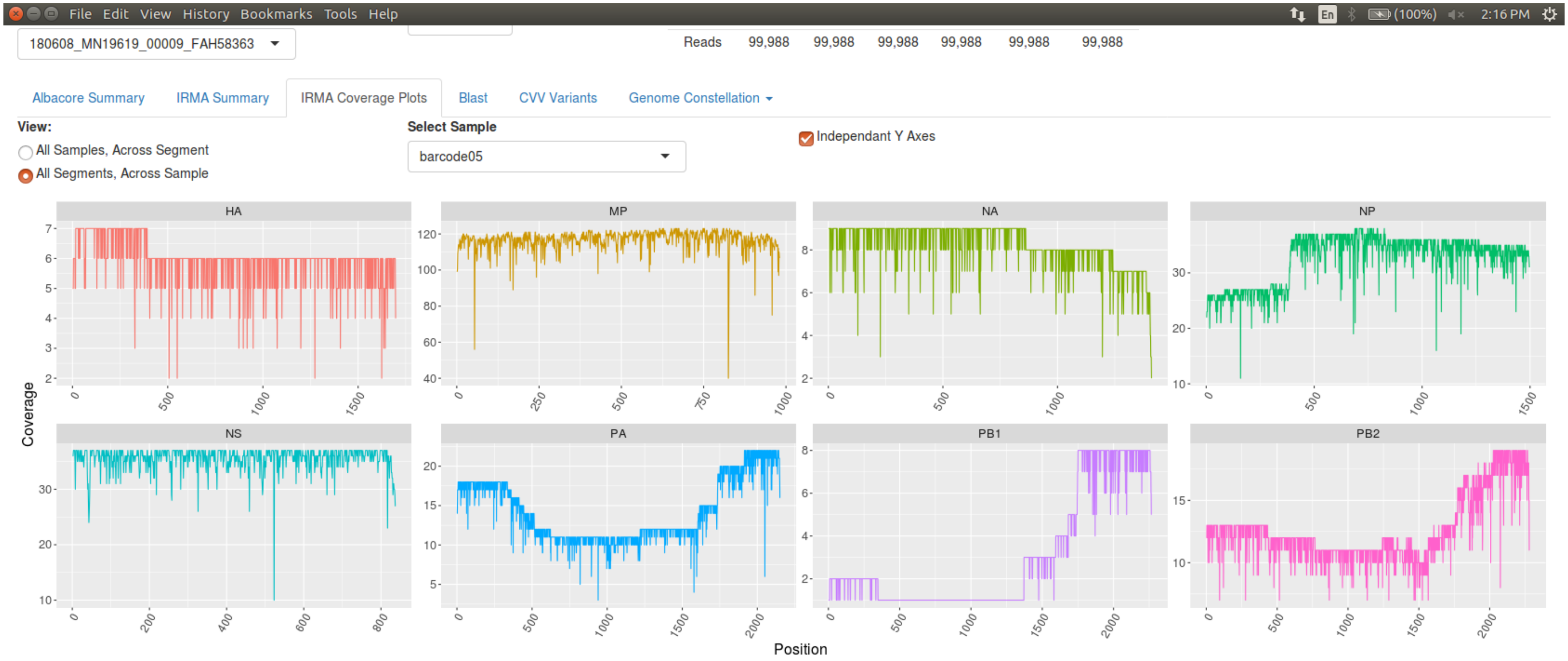


BC05 coverage screenshot

**Figure S4**

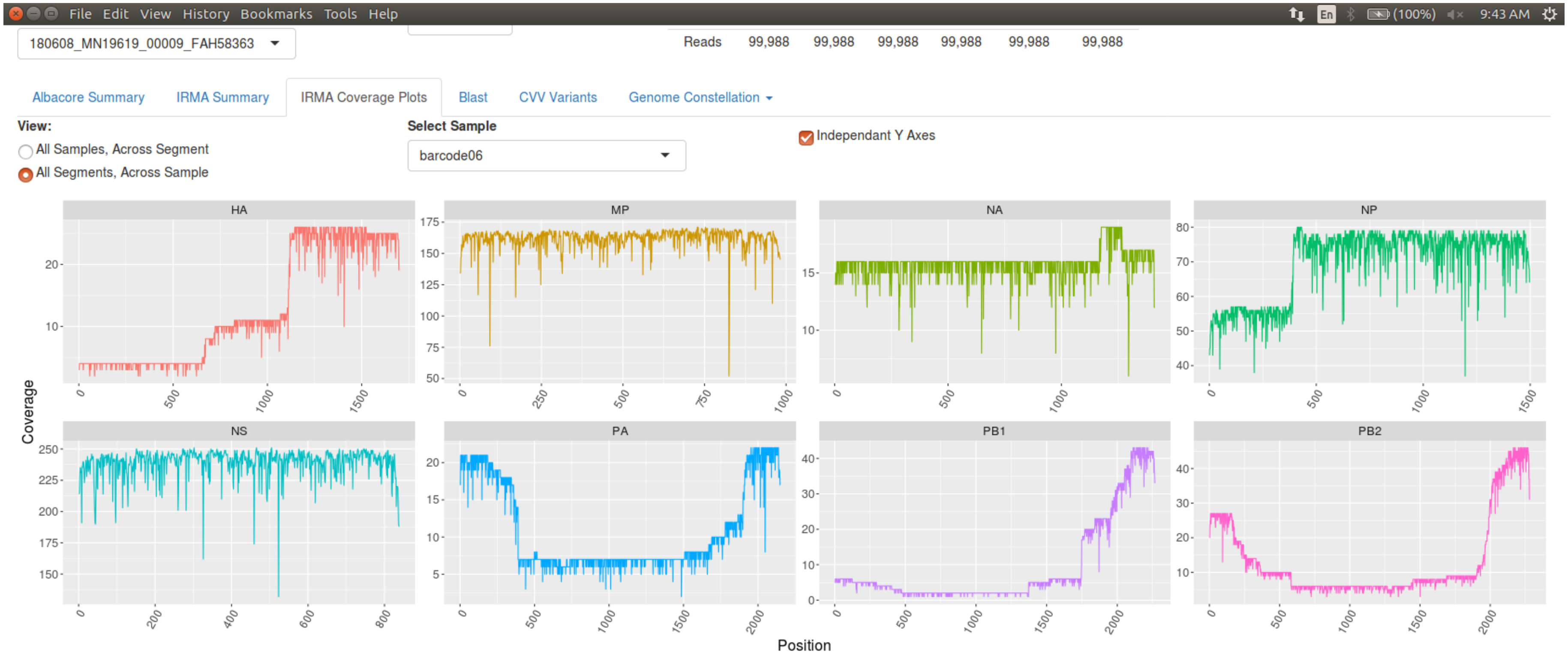


BC06 coverage screenshot

### Figure S5

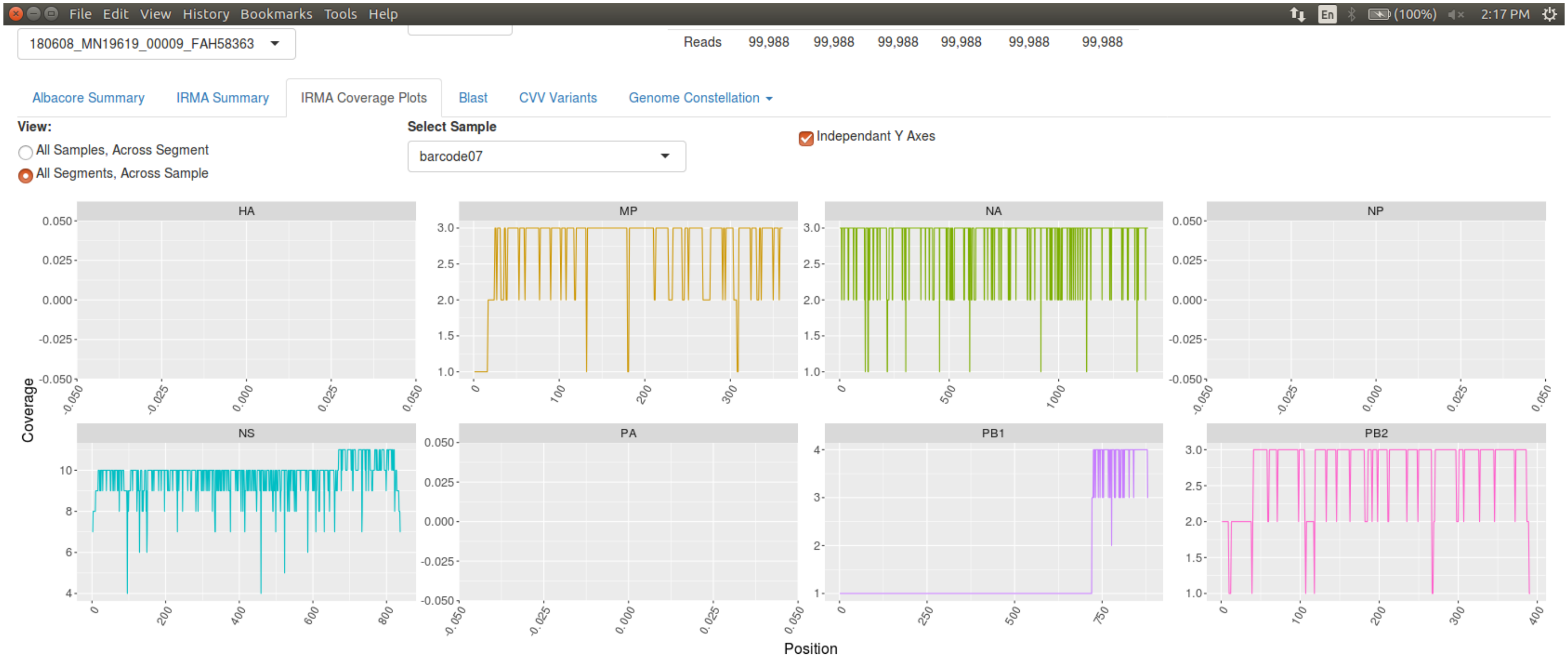


BC07 coverage screenshot

### Figure S6

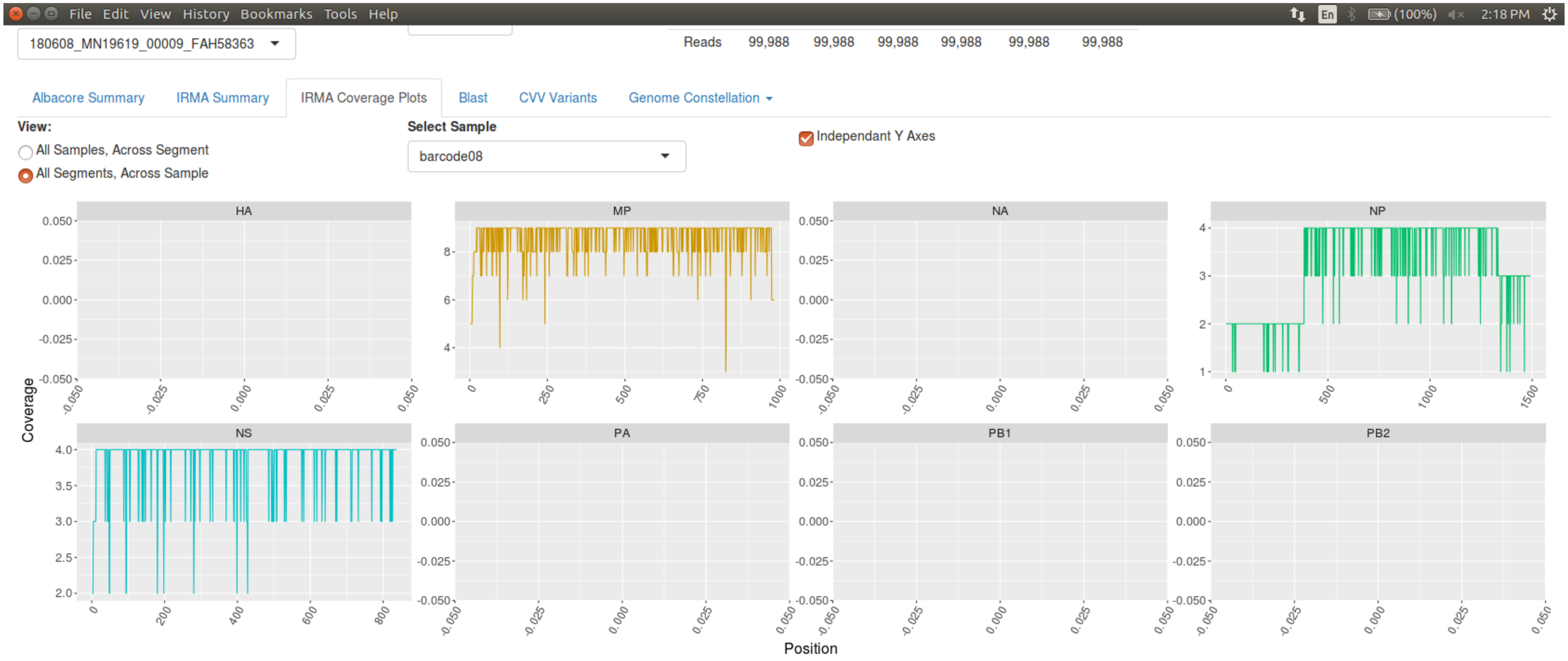


BC08 coverage screenshot

**Figure S7**

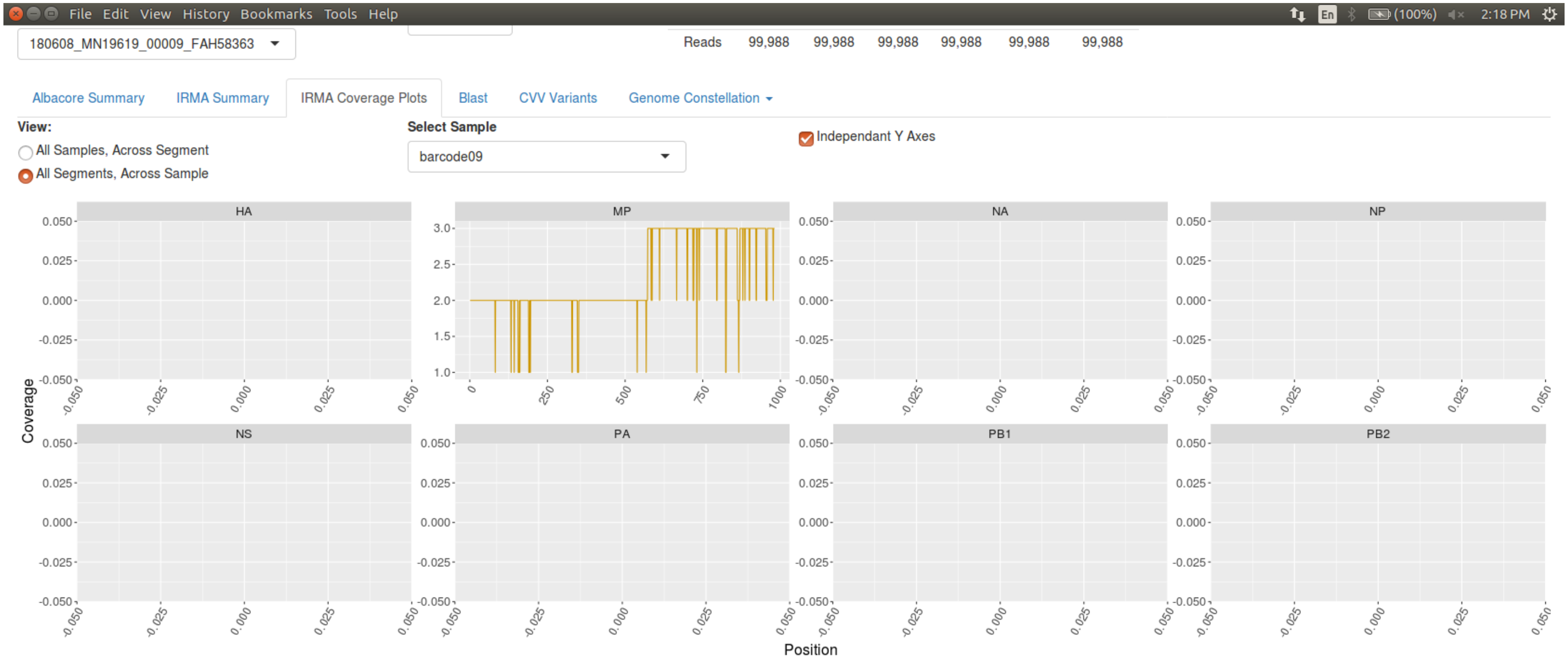


BC09 coverage screenshot

### Figure S8

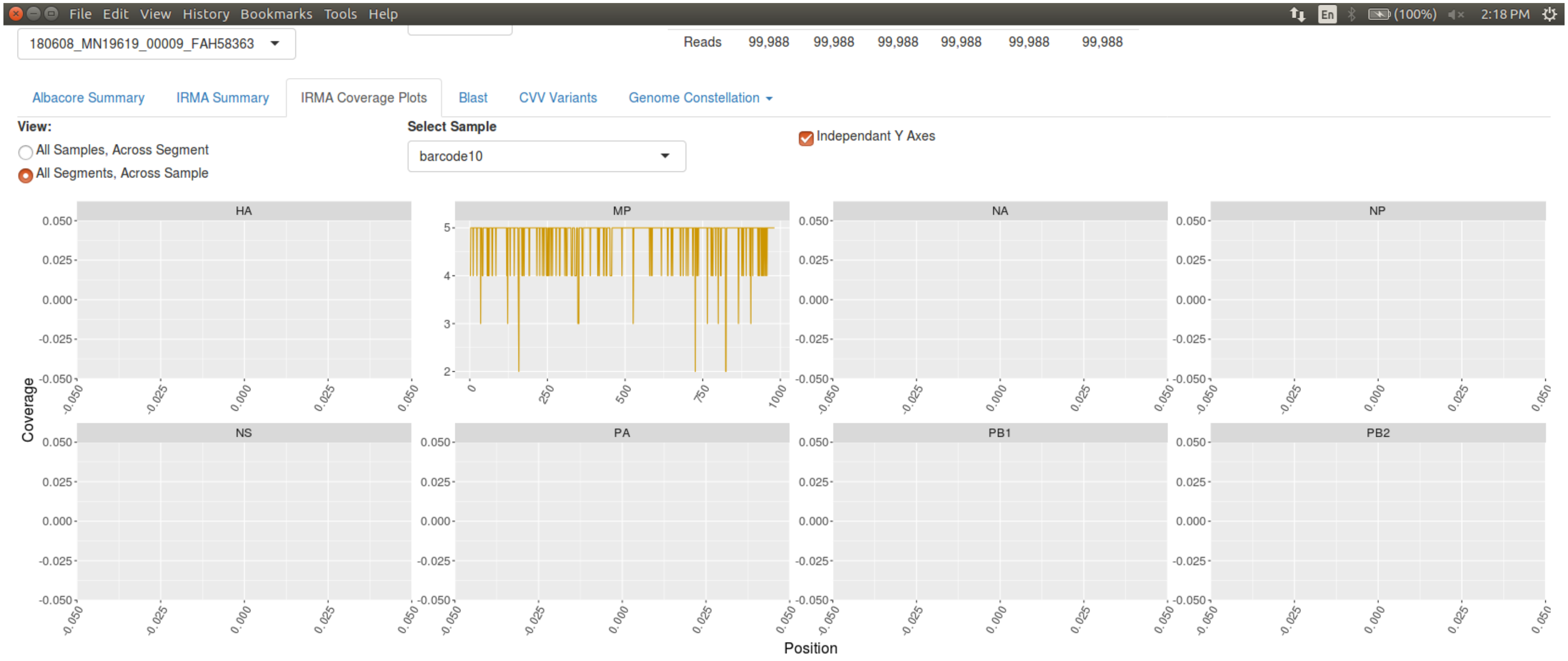


BC10 coverage screenshot

**Figure S9**

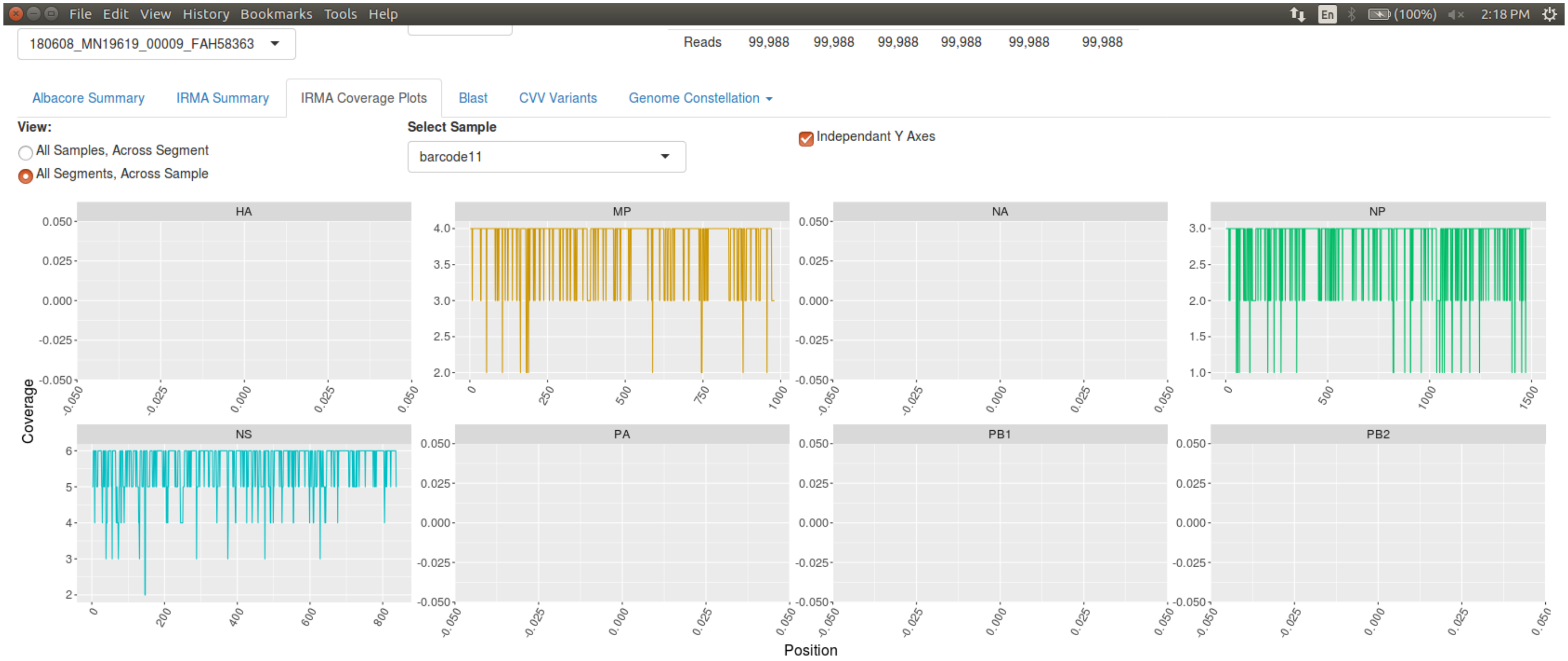


BC11 coverage screenshot

**Figure S10**

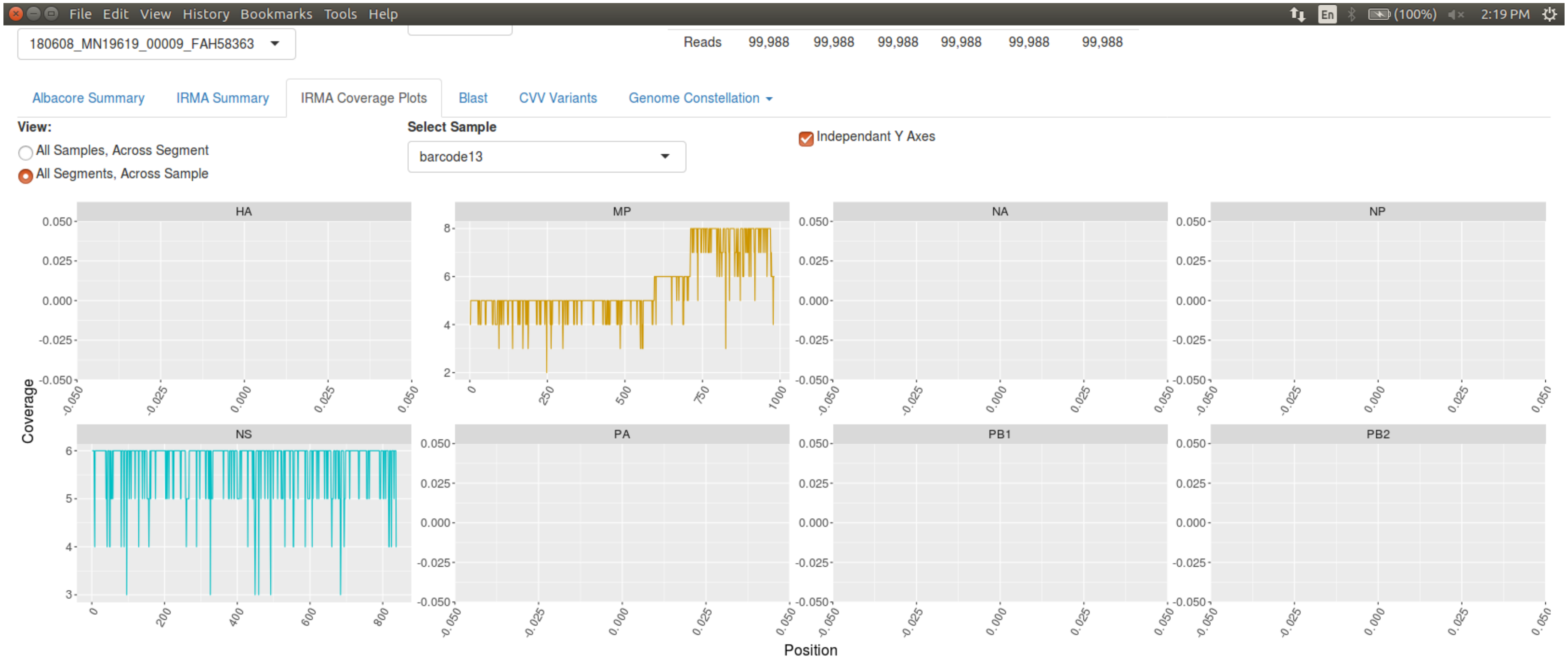


BC13 coverage screenshot

### Figure S11

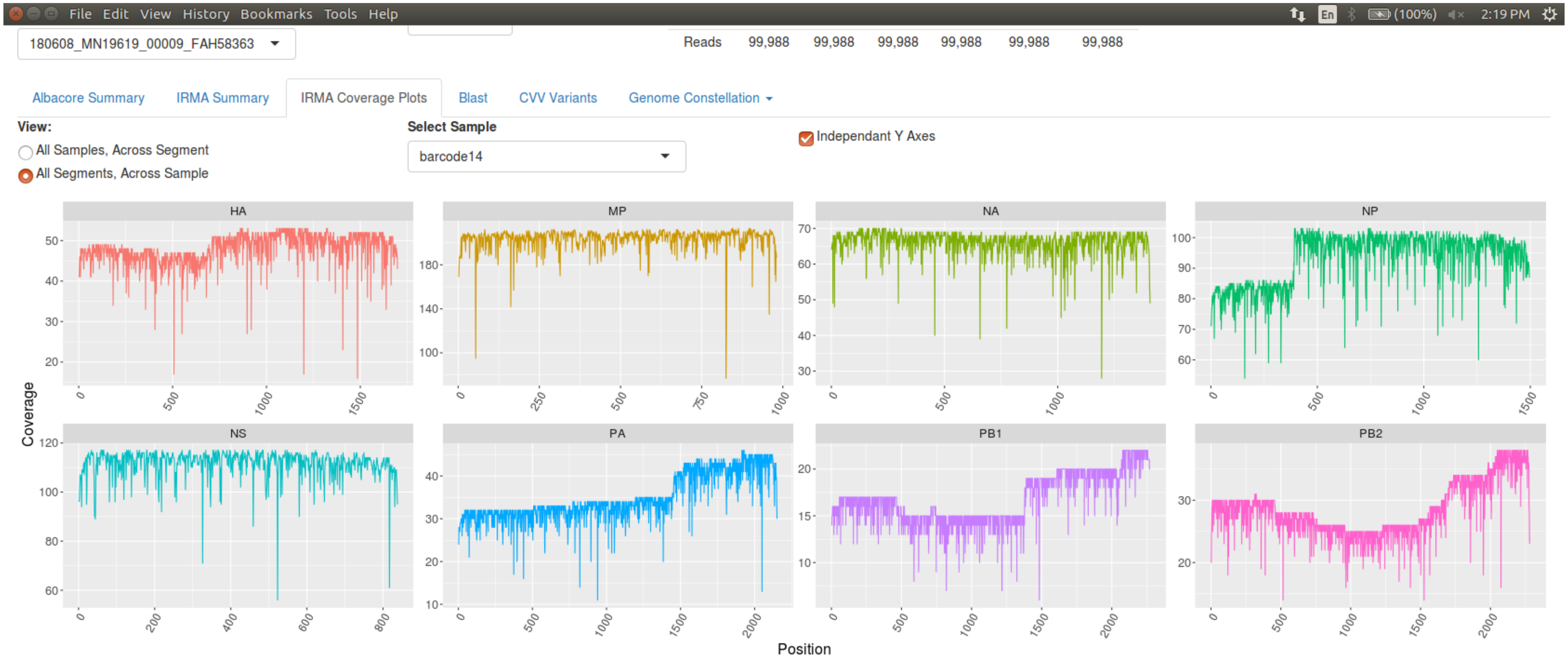


BC14 coverage screenshot

**Figure S12**

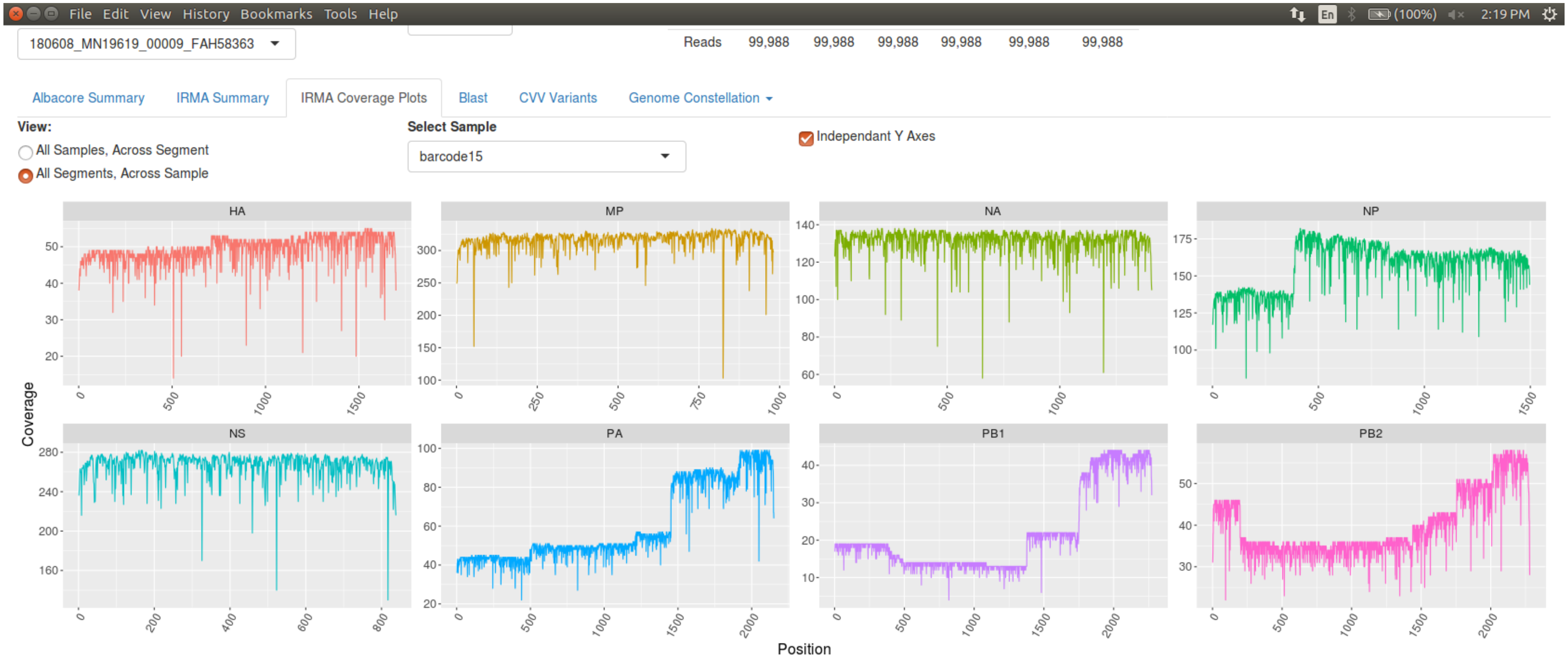


BC15 coverage screenshot

### Figure S13

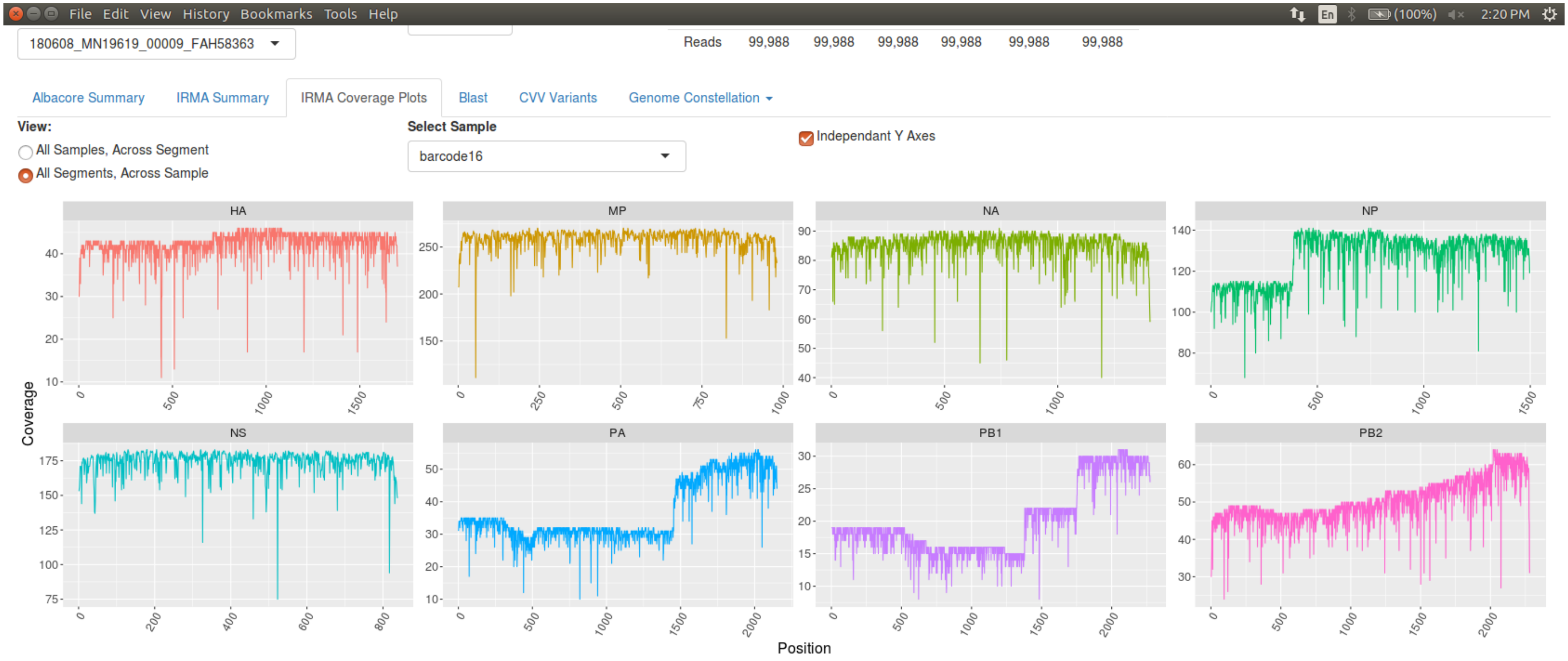


BC16 coverage screenshot

### Figure S14

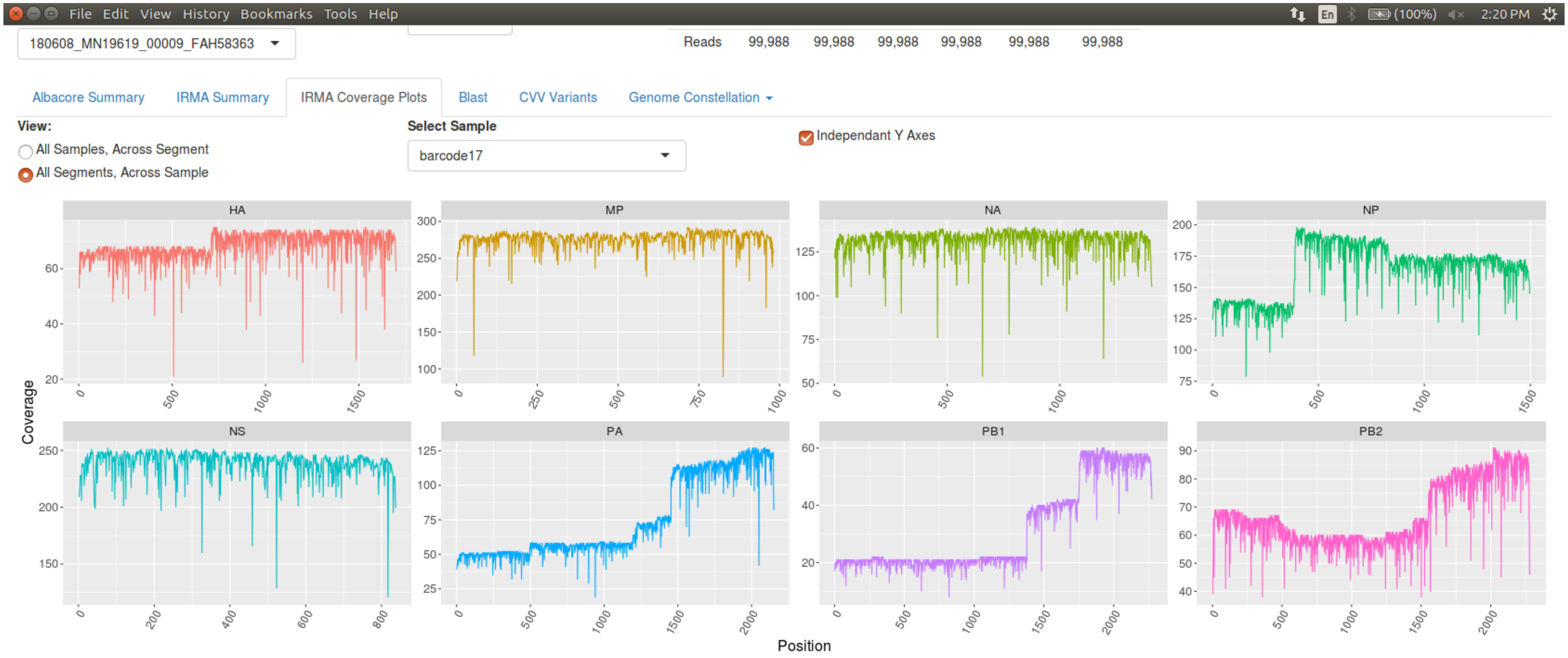


BC17 coverage screenshot

### Figure S15

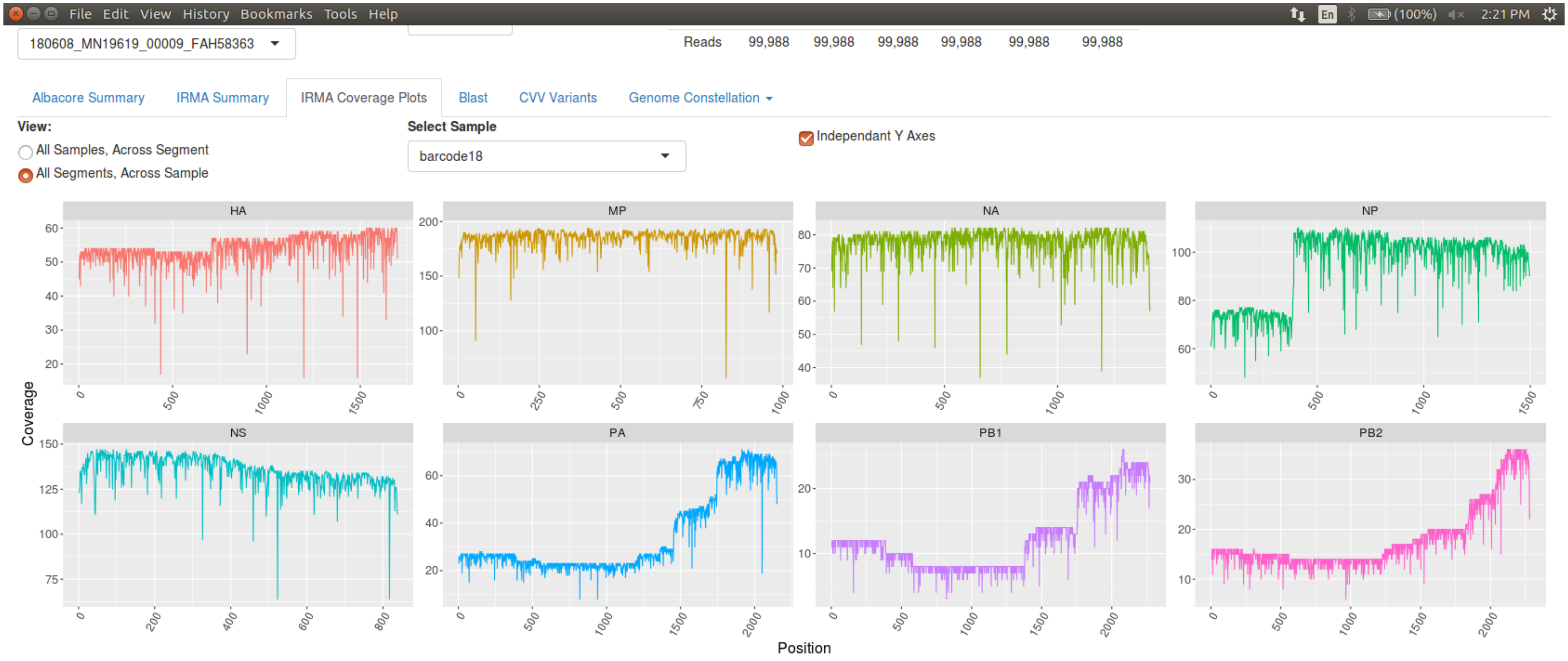


BC18 coverage screenshot

### Figure S16

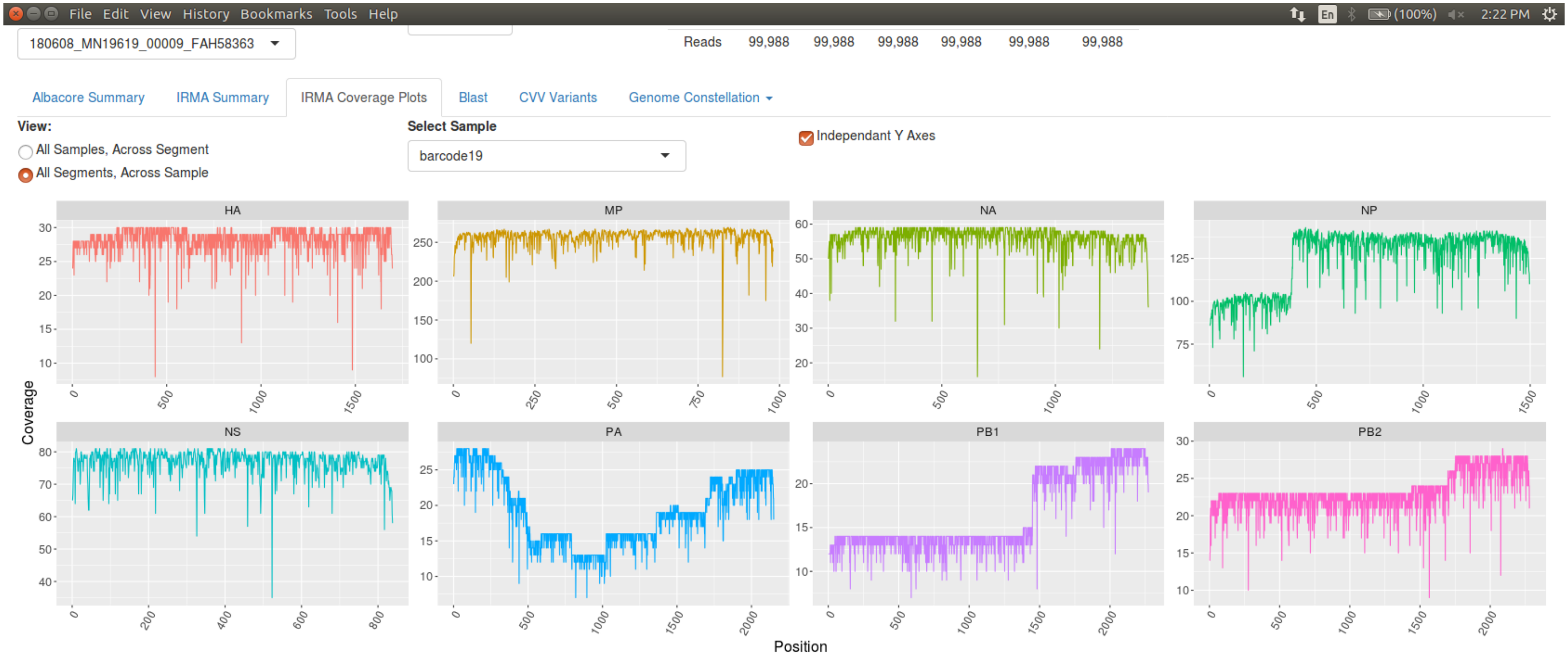


BC19 coverage screenshot

### Figure S17

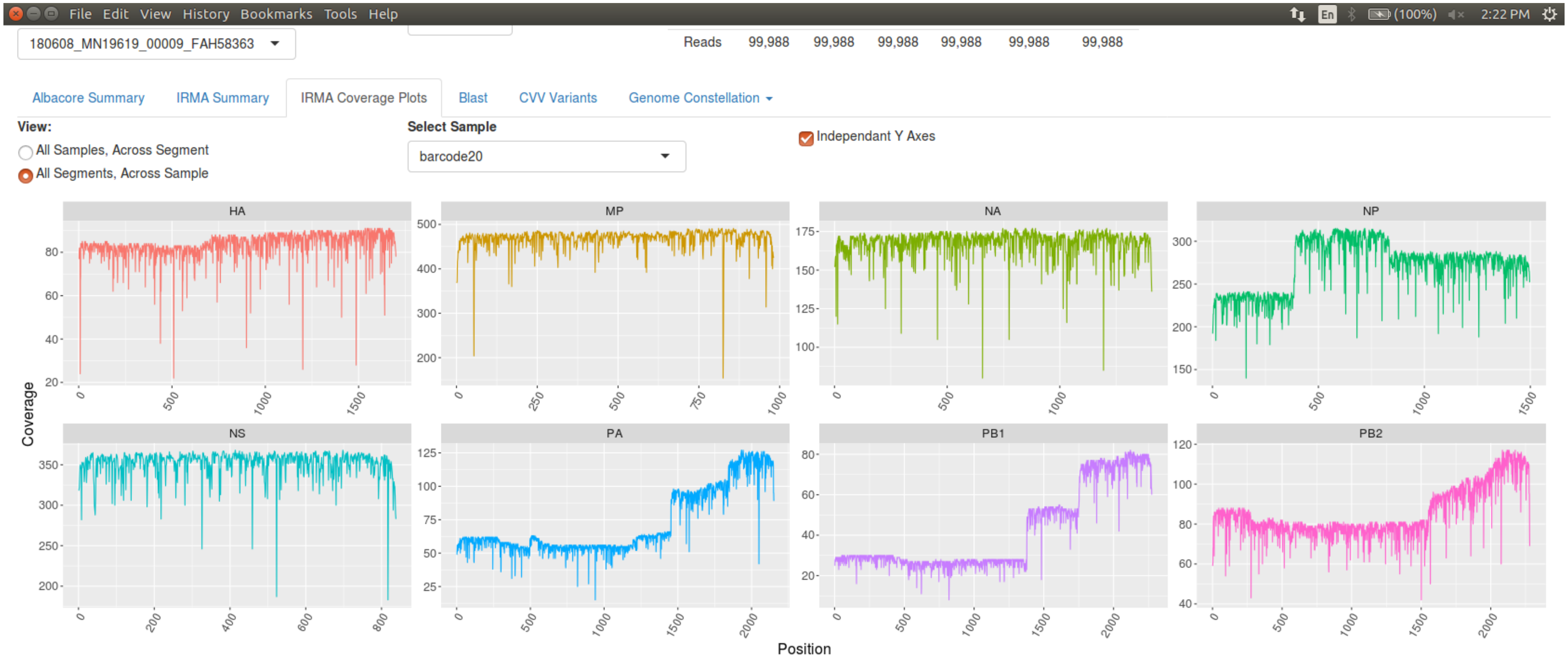


BC20 coverage screenshot

**Figure S18**

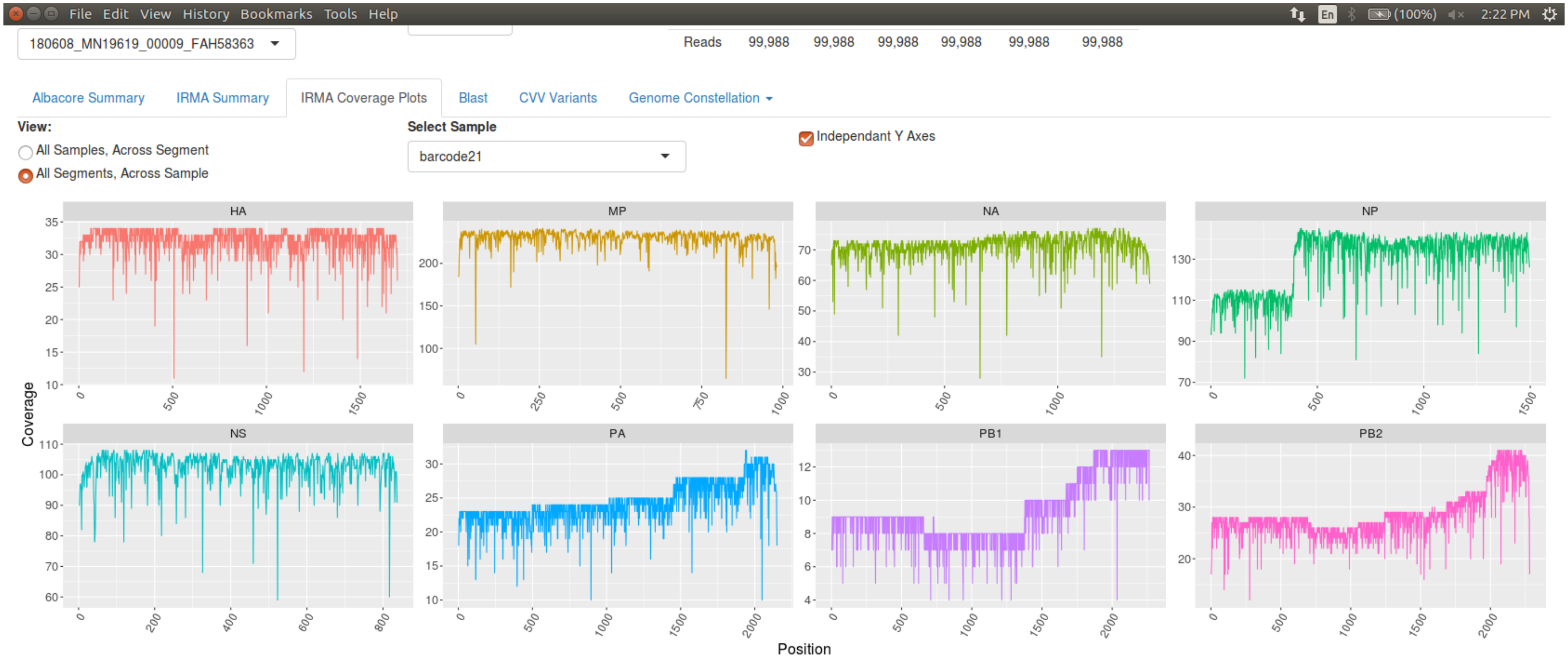


BC21 coverage screenshot

### Figure S19

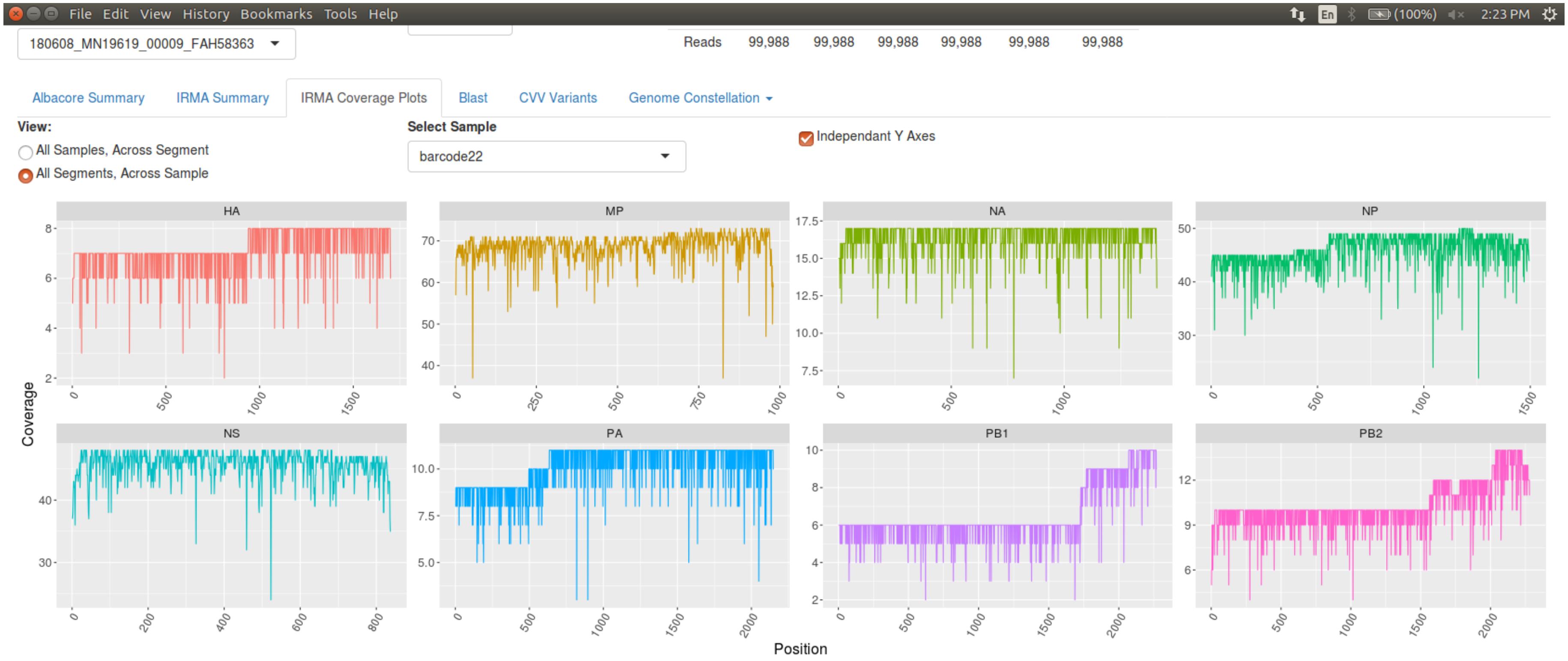


BC22 coverage screenshot

### Figure S20

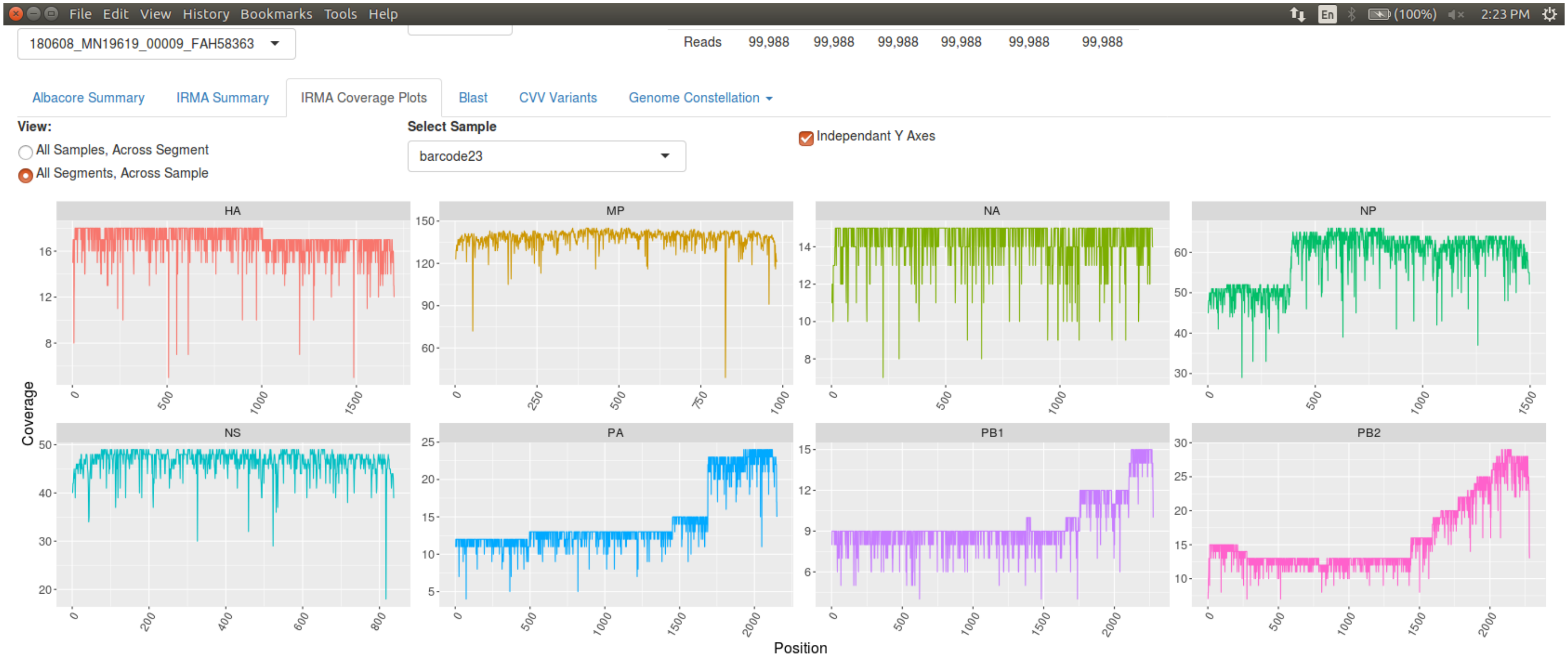


BC23 coverage diagram

### Figure S21

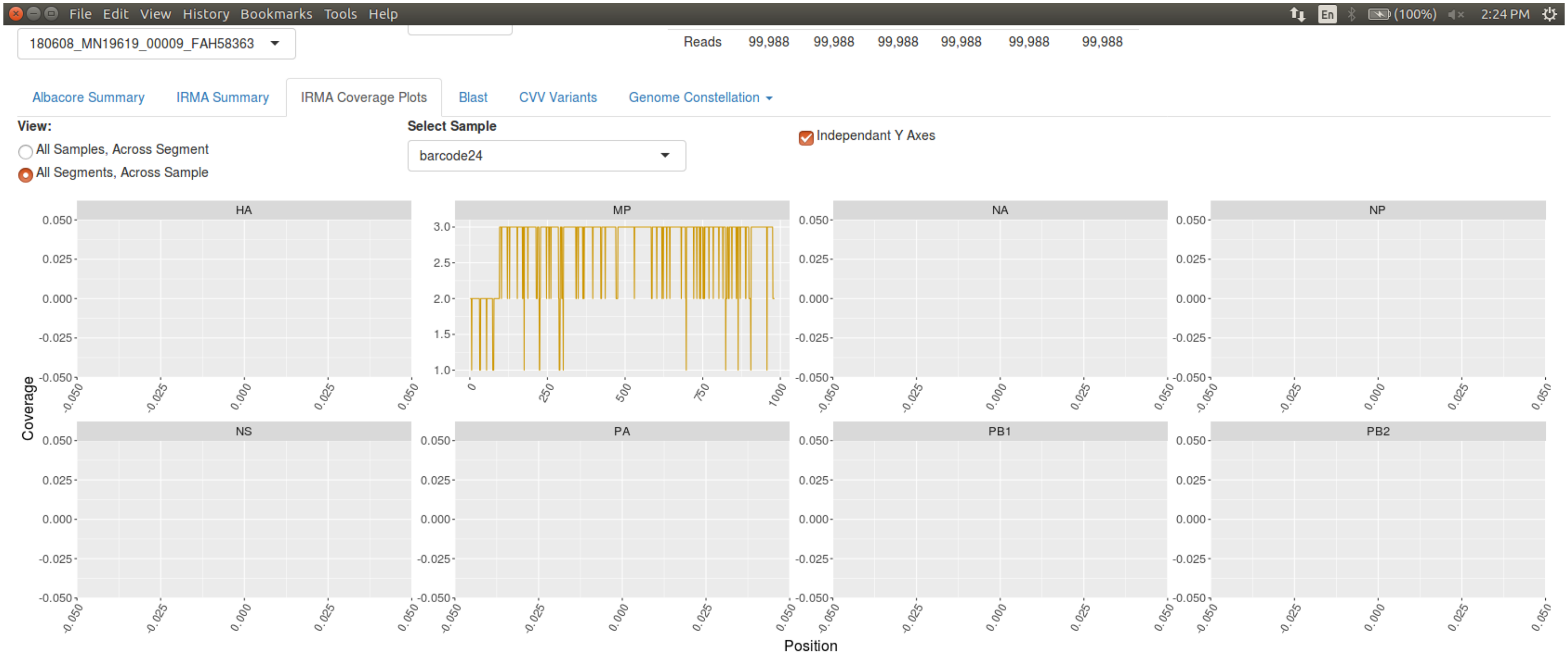


BC24 coverage screenshot

**Figure S22**

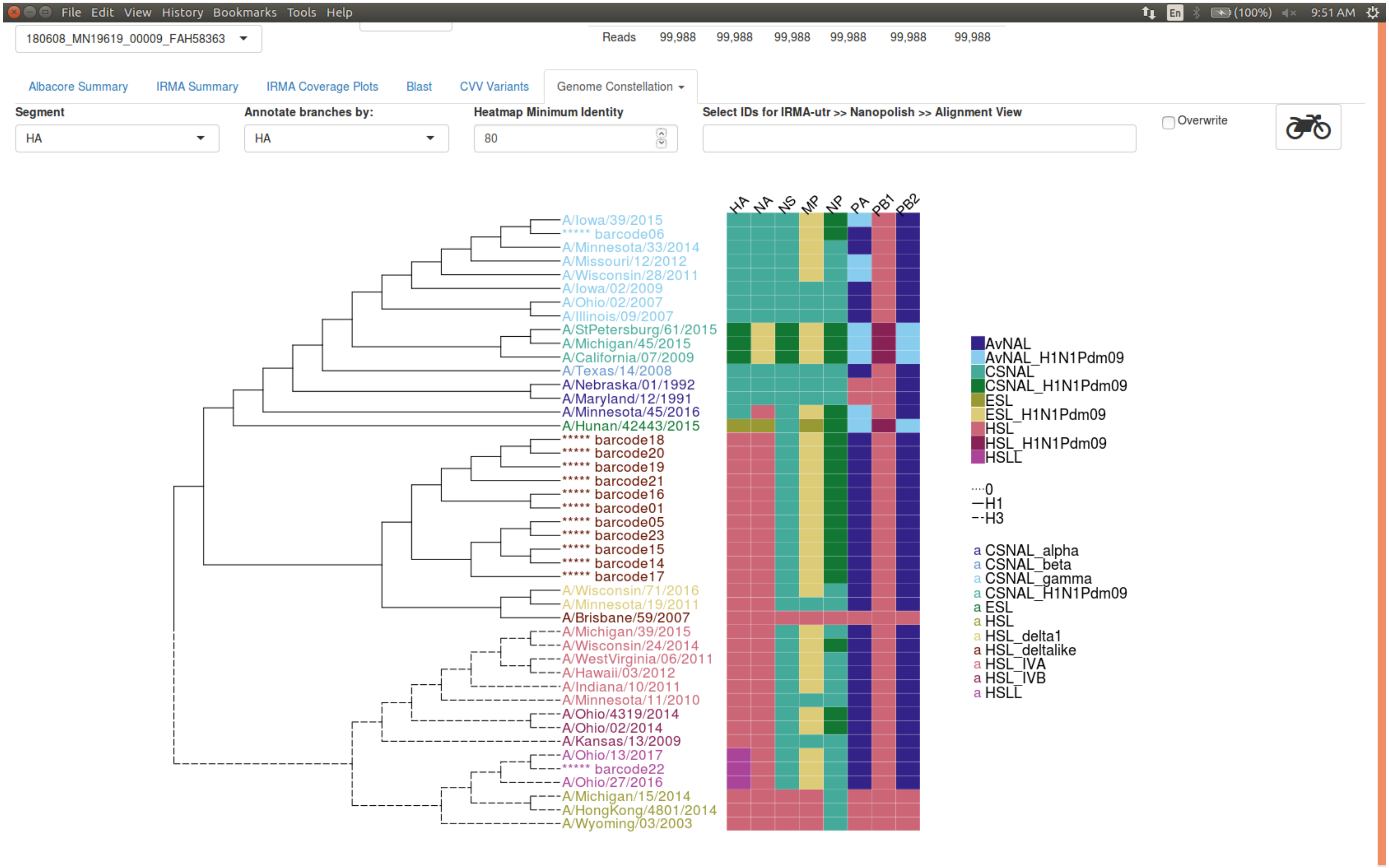
File Edit View History Bookmarks Tools Help
180608_MN19619_00009_FAH58363
Reads 99,988 99,988 99,988 99,988 99,988 99,988
Albacore Summary
IRMA Summary
IRMA Coverage Plots
Blast
CVV Variants
Genome Constellation
Segment
HA
Annotate branches by:
HA
Heatmap Minimum Identity
80
Select IDs for IRMA-utr >> Nanopolish >> Alignment View
Overwrite
HA NA NS MP NP PA PB1 PB2
A/Iowa/39/2015
***** barcode06
A/Minnesota/33/2014
A/Missouri/12/2012
A/Wisconsin/28/2011
A/Iowa/02/2009
A/Ohio/02/2007
A/Illinois/09/2007
A/StPetersburg/61/2015
A/Michigan/45/2015
A/California/07/2009
A/Texas/14/2008
A/Nebraska/01/1992
A/Maryland/12/1991
A/Minnesota/45/2016
A/Hunan/42443/2015
***** barcode18
***** barcode20
***** barcode19
***** barcode21
***** barcode16
***** barcode01
***** barcode05
***** barcode23
***** barcode15
***** barcode14
***** barcode17
A/Wisconsin/71/2016
A/Minnesota/19/2011
A/Brisbane/59/2007
A/Michigan/39/2015
A/Wisconsin/24/2014
A/WestVirginia/06/2011
A/Hawaii/03/2012
A/Indiana/10/2011
A/Minnesota/11/2010
A/Ohio/4319/2014
A/Ohio/02/2014
A/Kansas/13/2009
A/Ohio/13/2017
***** barcode22
A/Ohio/27/2016
A/Michigan/15/2014
A/HongKong/4801/2014
A/Wyoming/03/2003
AvNAL
AvNAL_H1N1Pdm09
CSNAL
CSNAL_H1N1Pdm09
ESL
ESL_H1N1Pdm09
HSL
HSL_H1N1Pdm09
HSLL
0
H1
H3
a CSNAL_alpha
a CSNAL_beta
a CSNAL_gamma
a CSNAL_H1N1Pdm09
a ESL
a HSL
a HSL_delta1
a HSL_deltalike
a HSL_IVA
a HSL_IVB
a HSLL

HA tree screenshot. Heatmap indicates the top Blast hit for a given sample's segment, over a user-defined minimum identity. Annotations are those used by the Zoonotic Virus Team at CDC and include: Avian North American Lineage (AvNAL), Avian North American 2009 H1N1 Pandemic Lineage (AvNAL_H1N1Pdm09), Classical Swine North American Lineage (CSNAL), Classical Swine North American 2009 H1N1 Pandemic Lineage (CSNAL_H1N1Pdm09), European Swine Lineage (ESL), European Swine 2009 H1N1 Pandemic Lineage (ESL_H1N1Pdm09), Human Seasonal Lineage (HSL), Human Seasonal 2009 H1N1 Pandemic Lineage (HSL_H1N1Pdm09) and Human Seasonal-Like Lineage (HSLL). Further HA annotations are viewed by the coloring of strain names and include: alpha, beta, gamma and H1N1pdm09 subclades of Classical Swine North American Lineages, delta-1 and delta-like (more contemporaneously referred to as delta-2) subclades of Human Seasonal H1 Lineage and IVA and IVB subclades of Human Seasonal H3 Lineage.

**Figure S23**

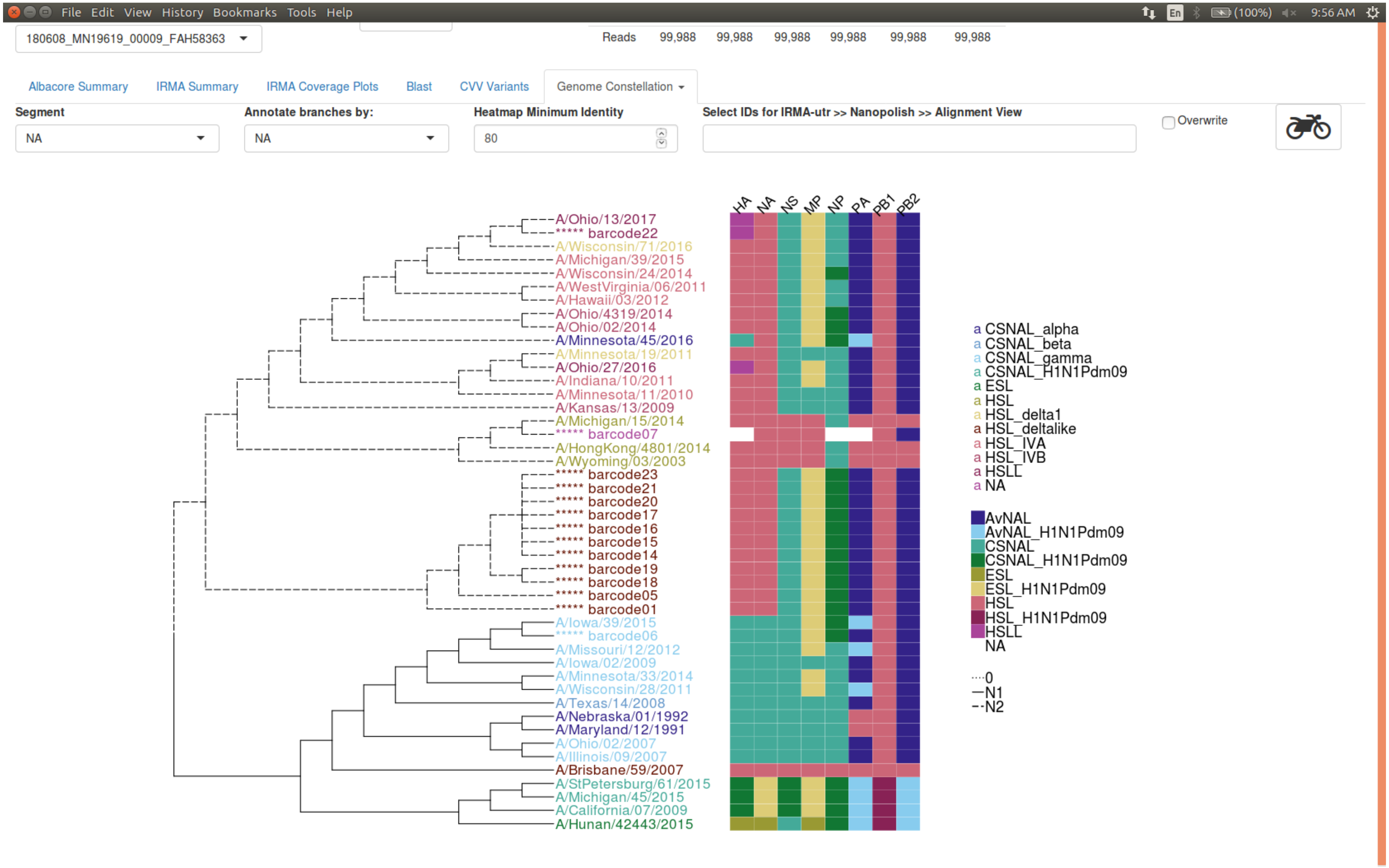


NA tree screenshot. Annotations are described in Figure S22.

### Figure S24

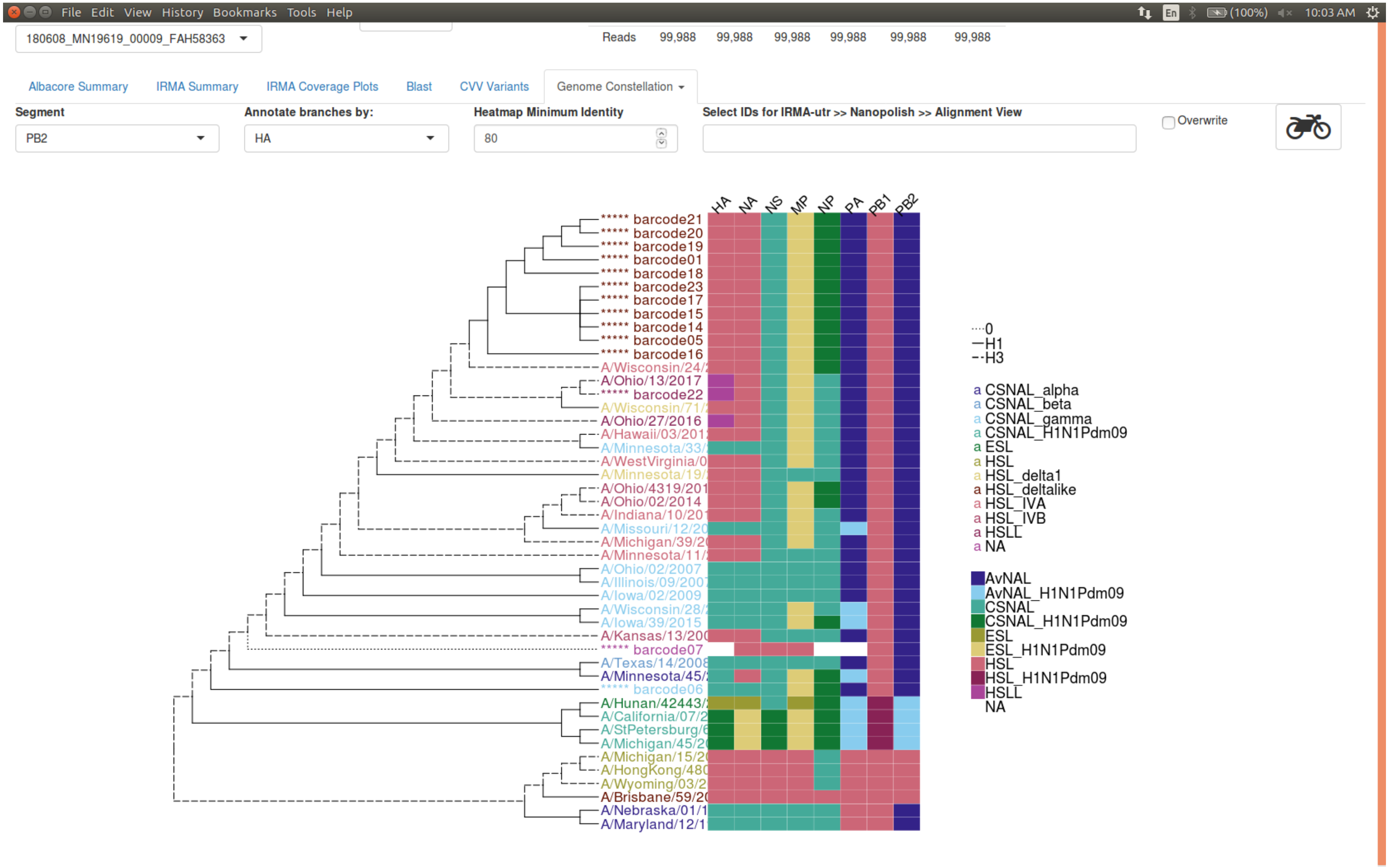

PB2 tree screenshot. Annotations are described in Figure S22.

### Figure S25

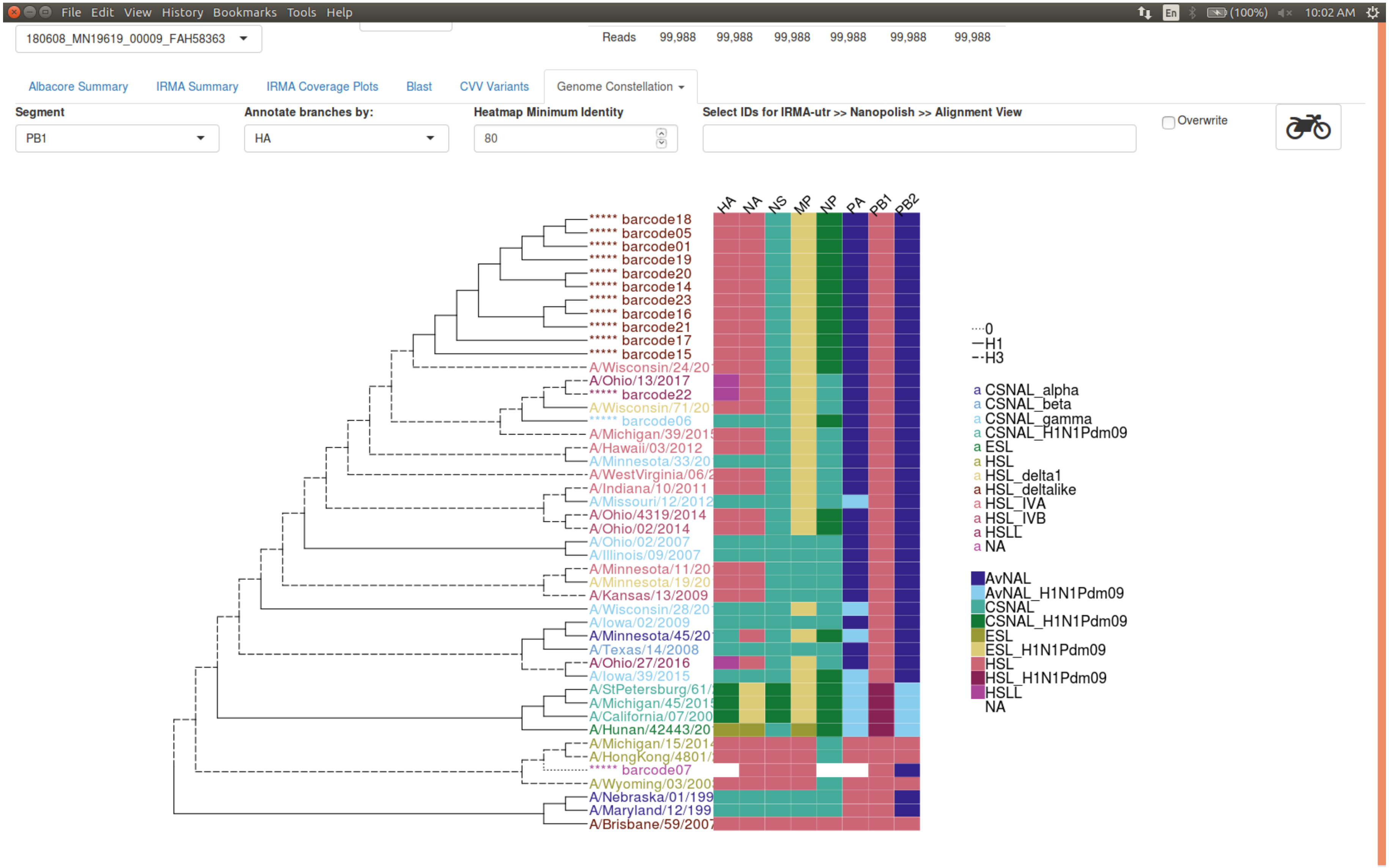


PB1 tree screenshot. Annotations are described in Figure S22.

### Figure S26

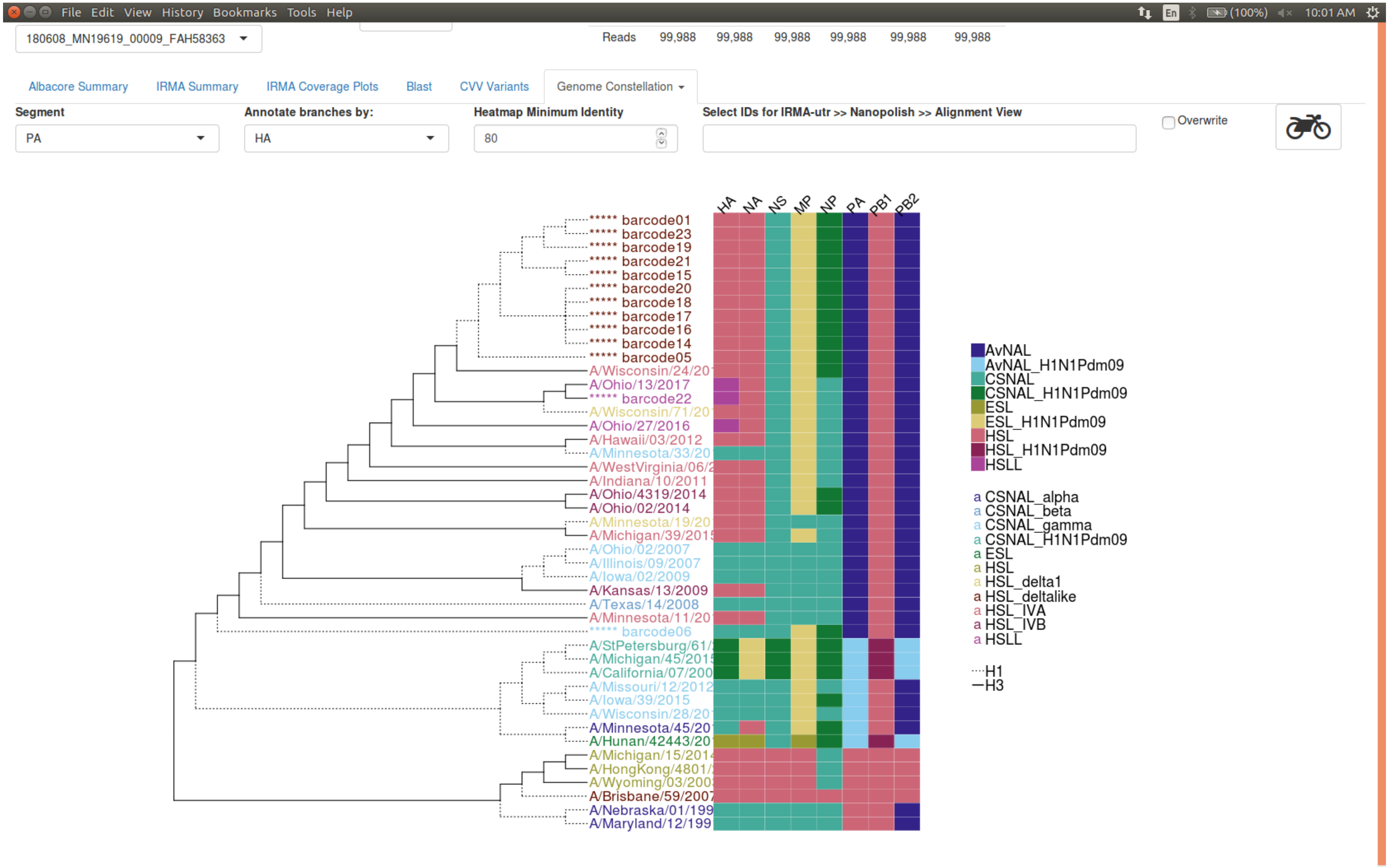


PA tree screenshot. Annotations are described in Figure S22.

### Figure S27

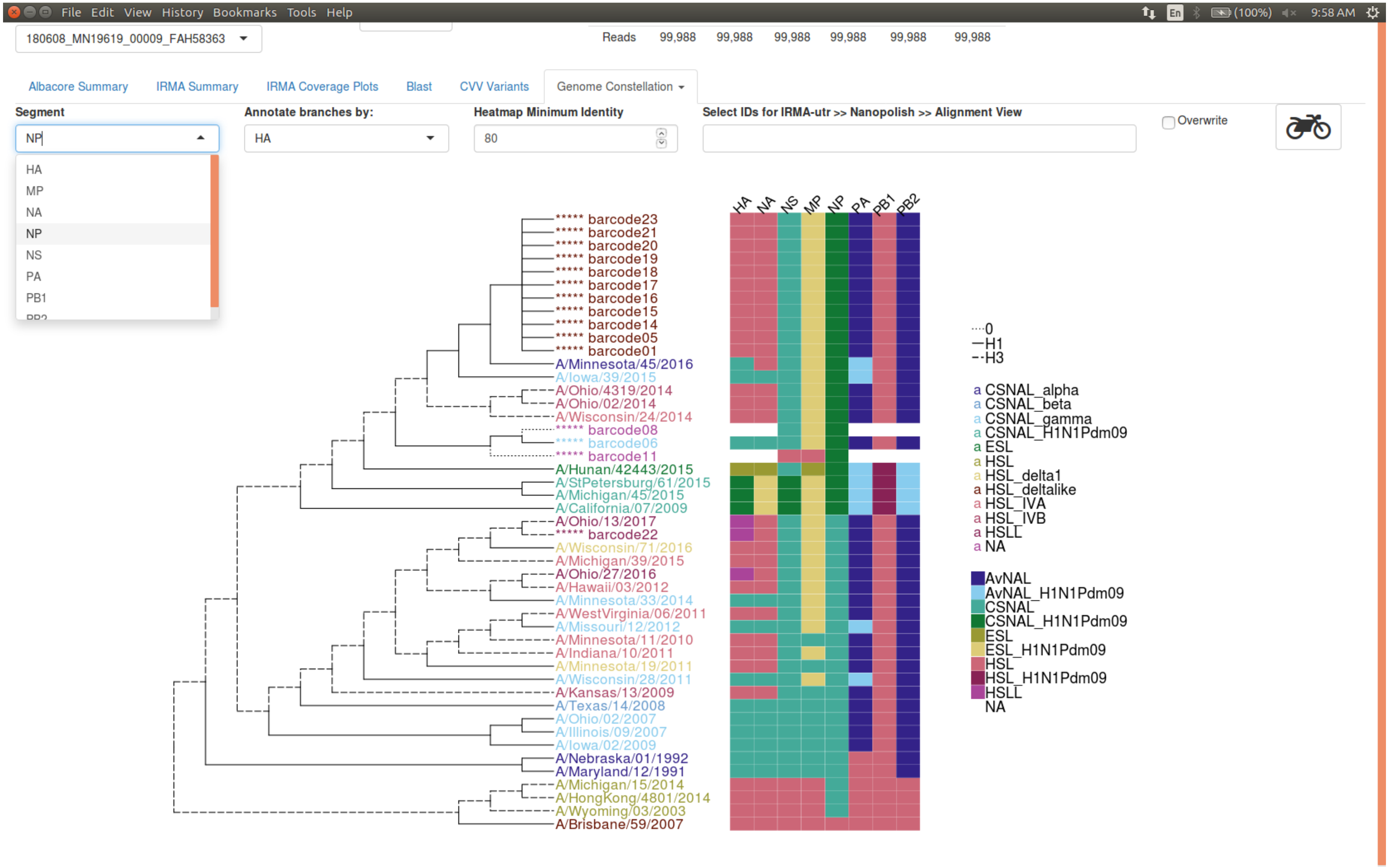


NP tree screenshot. Annotations are described in Figure S22.

**Figure S28**

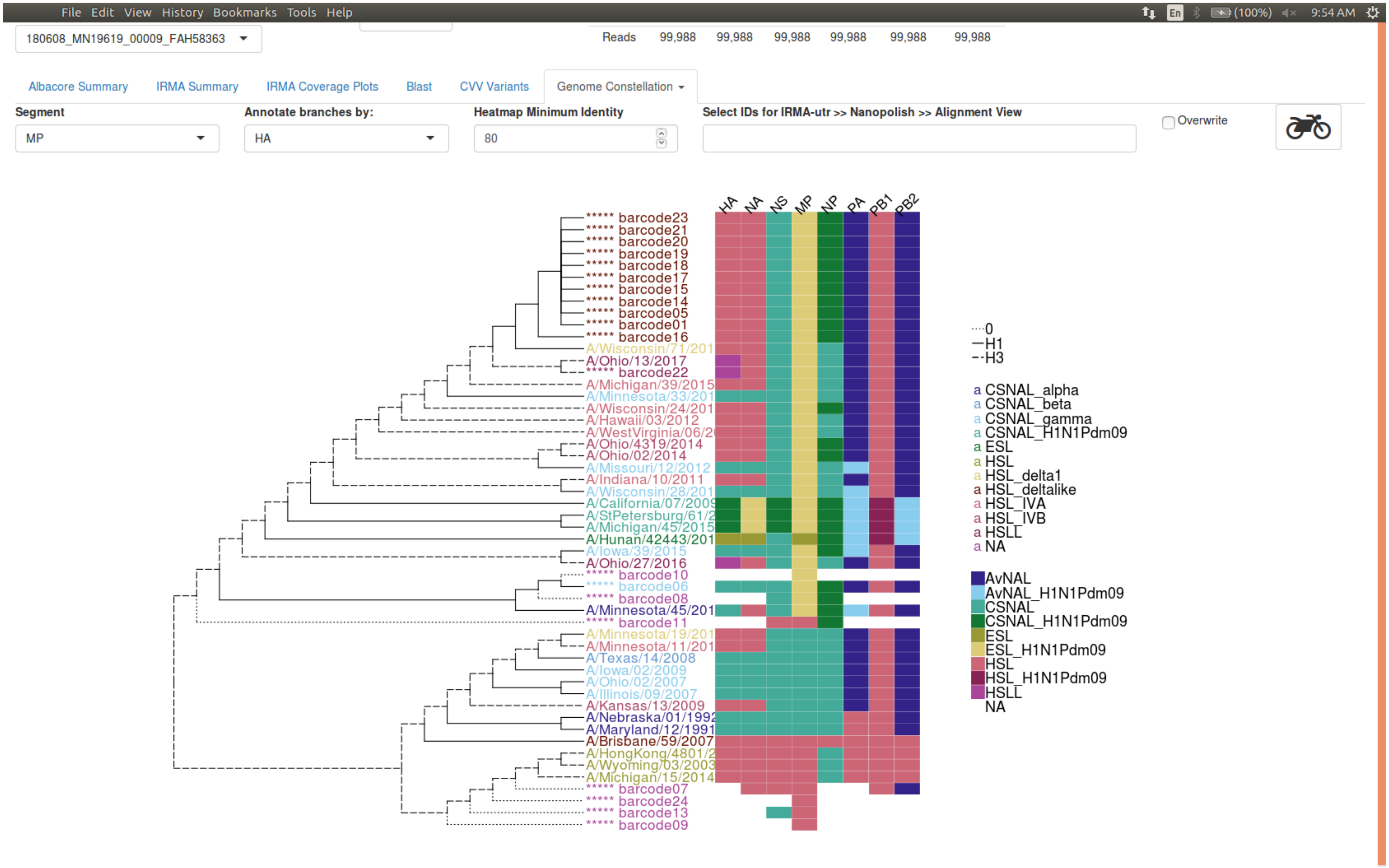


M tree screenshot. Annotations are described in Figure S22.

**Figure S29**

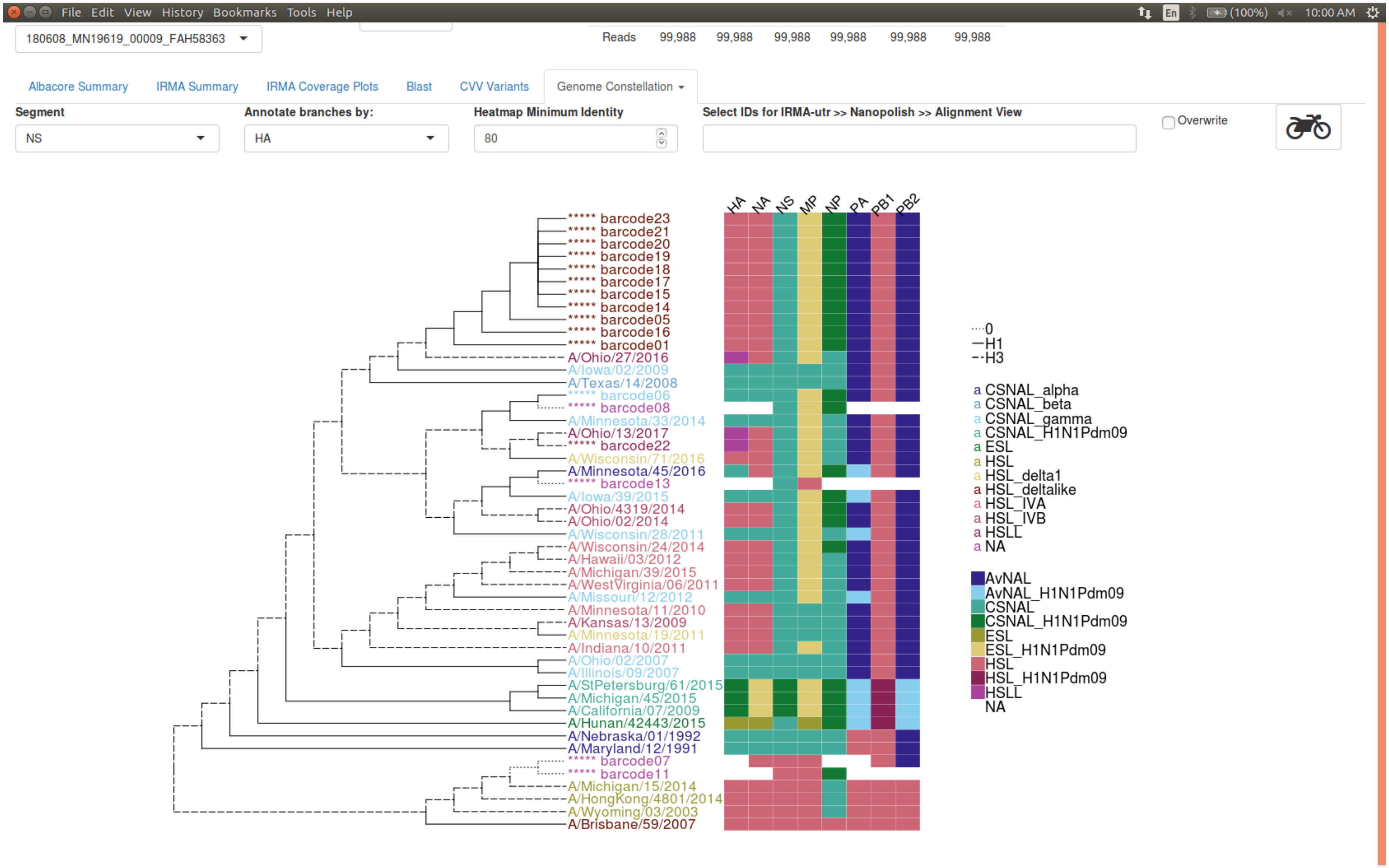


NS tree screenshot. Annotations are described in Figure S22.

### Figure S30

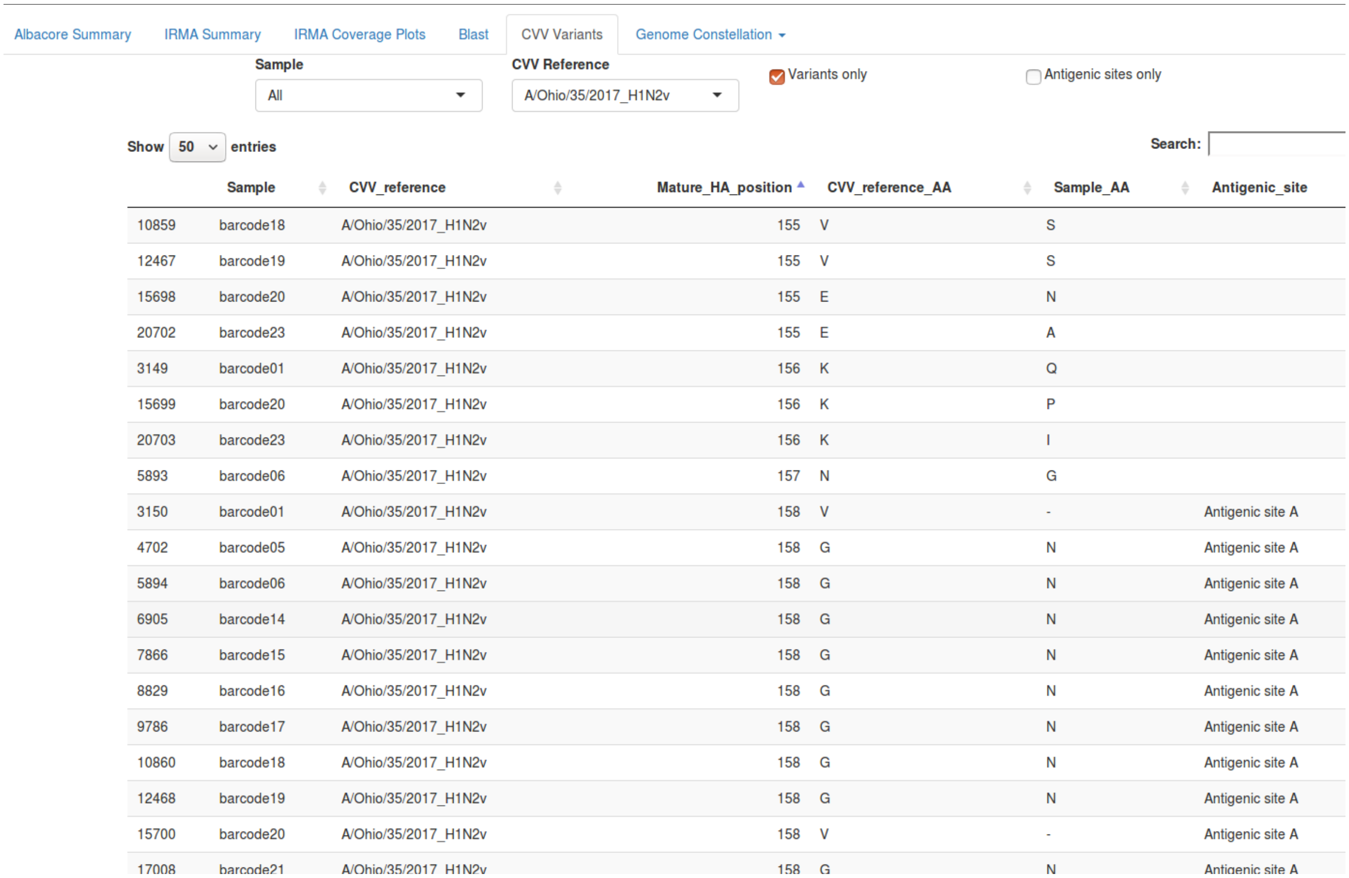


| | Sample | CVV_reference | Mature_HA_position | CVV_reference_AA | Sample_AA | Antigenic_site |
|---|---|---|---|---|---|---|
| 10859 | barcode18 | A/Ohio/35/2017_H1N2v | 155 | V | S | |
| 12467 | barcode19 | A/Ohio/35/2017_H1N2v | 155 | V | S | |
| 15698 | barcode20 | A/Ohio/35/2017_H1N2v | 155 | E | N | |
| 20702 | barcode23 | A/Ohio/35/2017_H1N2v | 155 | E | A | |
| 3149 | barcode01 | A/Ohio/35/2017_H1N2v | 156 | K | Q | |
| 15699 | barcode20 | A/Ohio/35/2017_H1N2v | 156 | K | P | |
| 20703 | barcode23 | A/Ohio/35/2017_H1N2v | 156 | K | I | |
| 5893 | barcode06 | A/Ohio/35/2017_H1N2v | 157 | N | G | |
| 3150 | barcode01 | A/Ohio/35/2017_H1N2v | 158 | V | - | Antigenic site A |
| 4702 | barcode05 | A/Ohio/35/2017_H1N2v | 158 | G | N | Antigenic site A |
| 5894 | barcode06 | A/Ohio/35/2017_H1N2v | 158 | G | N | Antigenic site A |
| 6905 | barcode14 | A/Ohio/35/2017_H1N2v | 158 | G | N | Antigenic site A |
| 7866 | barcode15 | A/Ohio/35/2017_H1N2v | 158 | G | N | Antigenic site A |
| 8829 | barcode16 | A/Ohio/35/2017_H1N2v | 158 | G | N | Antigenic site A |
| 9786 | barcode17 | A/Ohio/35/2017_H1N2v | 158 | G | N | Antigenic site A |
| 10860 | barcode18 | A/Ohio/35/2017_H1N2v | 158 | G | N | Antigenic site A |
| 12468 | barcode19 | A/Ohio/35/2017_H1N2v | 158 | G | N | Antigenic site A |
| 15700 | barcode20 | A/Ohio/35/2017_H1N2v | 158 | V | - | Antigenic site A |
| 17008 | barcode21 | A/Ohio/35/2017_H1N2v | 158 | G | N | Antigenic site A |

CVV Variants tab screenshot details numerous amino acid differences to current WHO Candidate Vaccine Viruses, including changes in known antigenic sites.

**Figure S31**

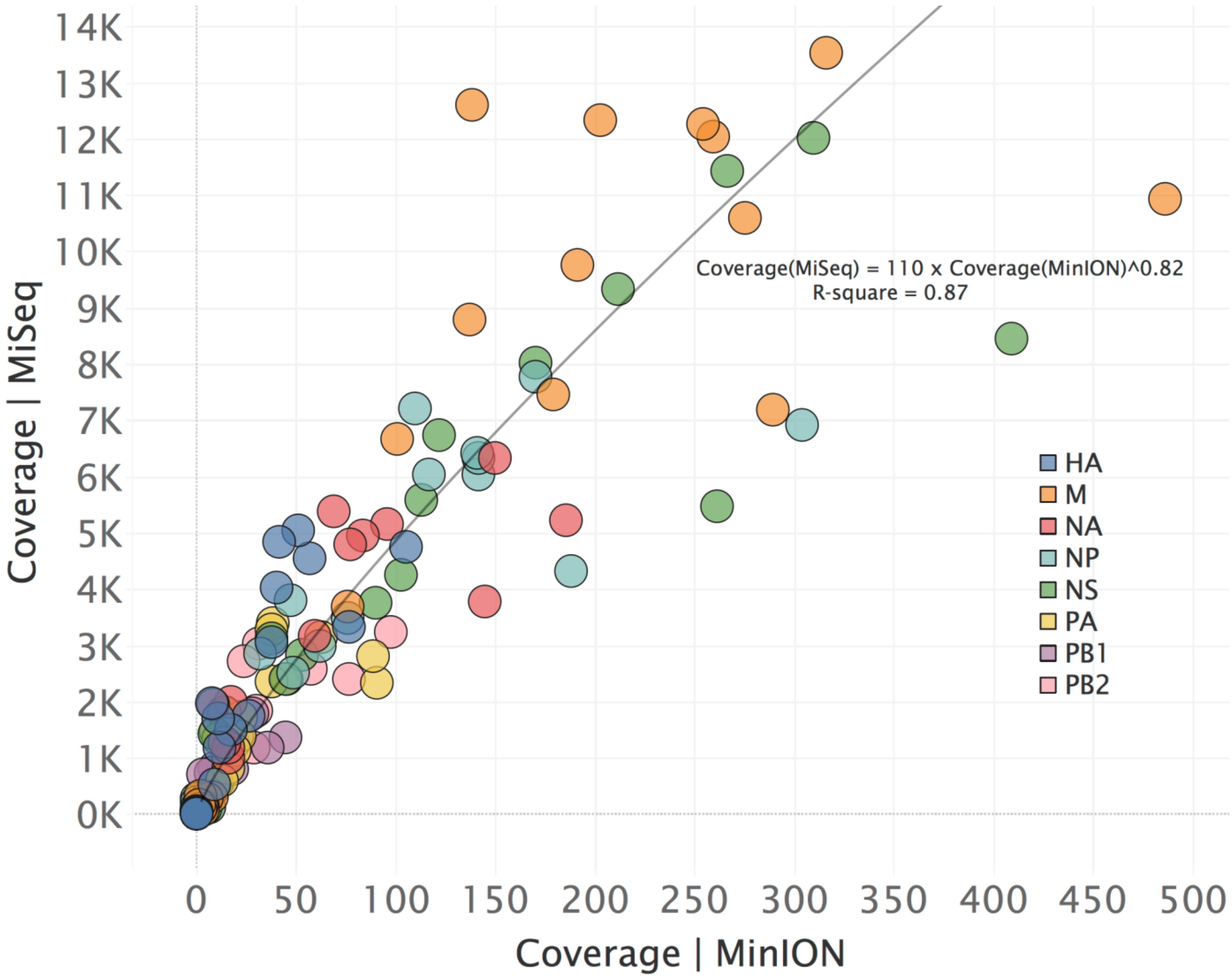


MinION vs MiSeq coverages are highly concordant, albeit on different scales.

**Figure S32**

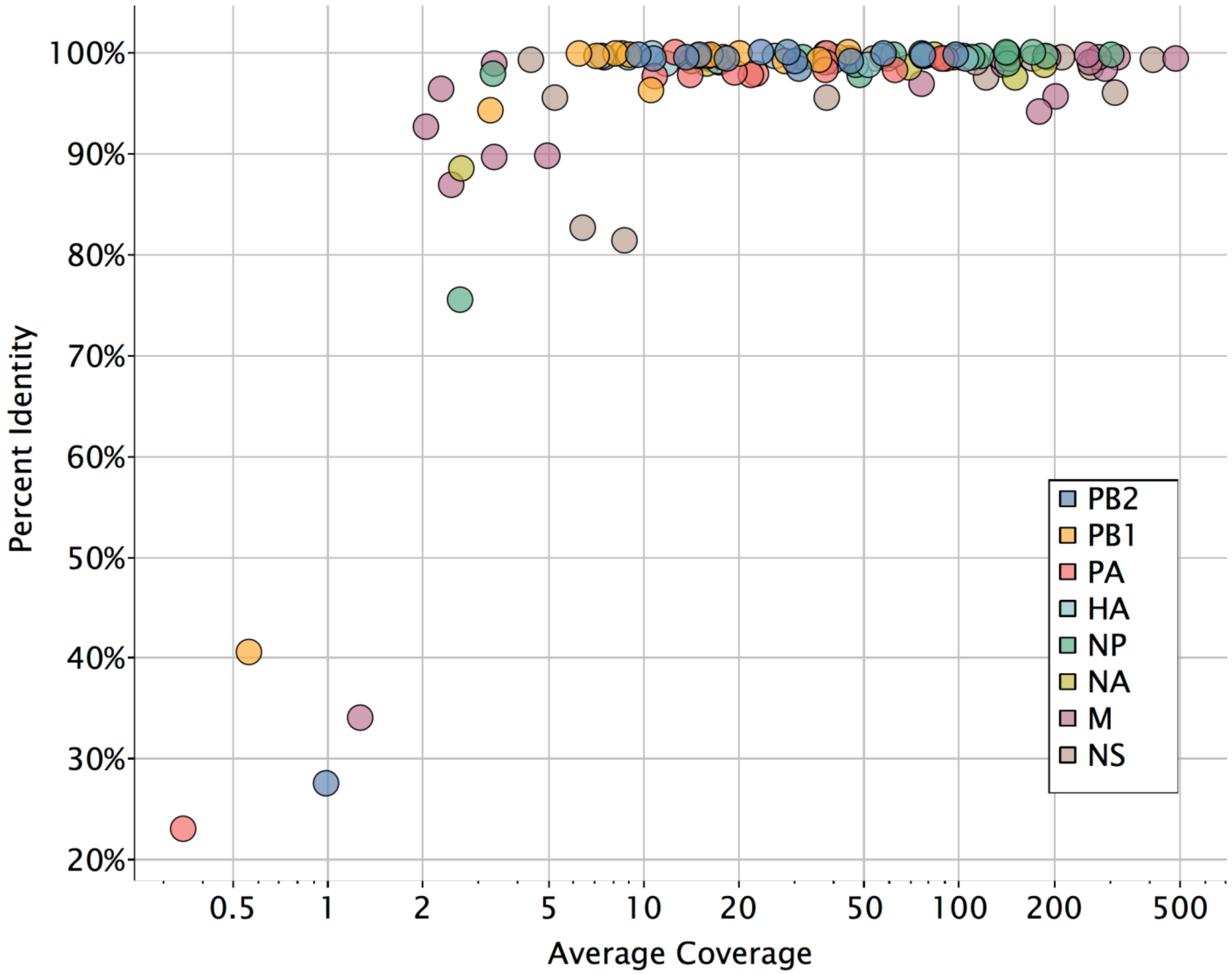


MinION coverage vs identity to MiSeq. MinION derived sequences achieve maximum consensus identity at 10-fold coverage.

**Figure S33**

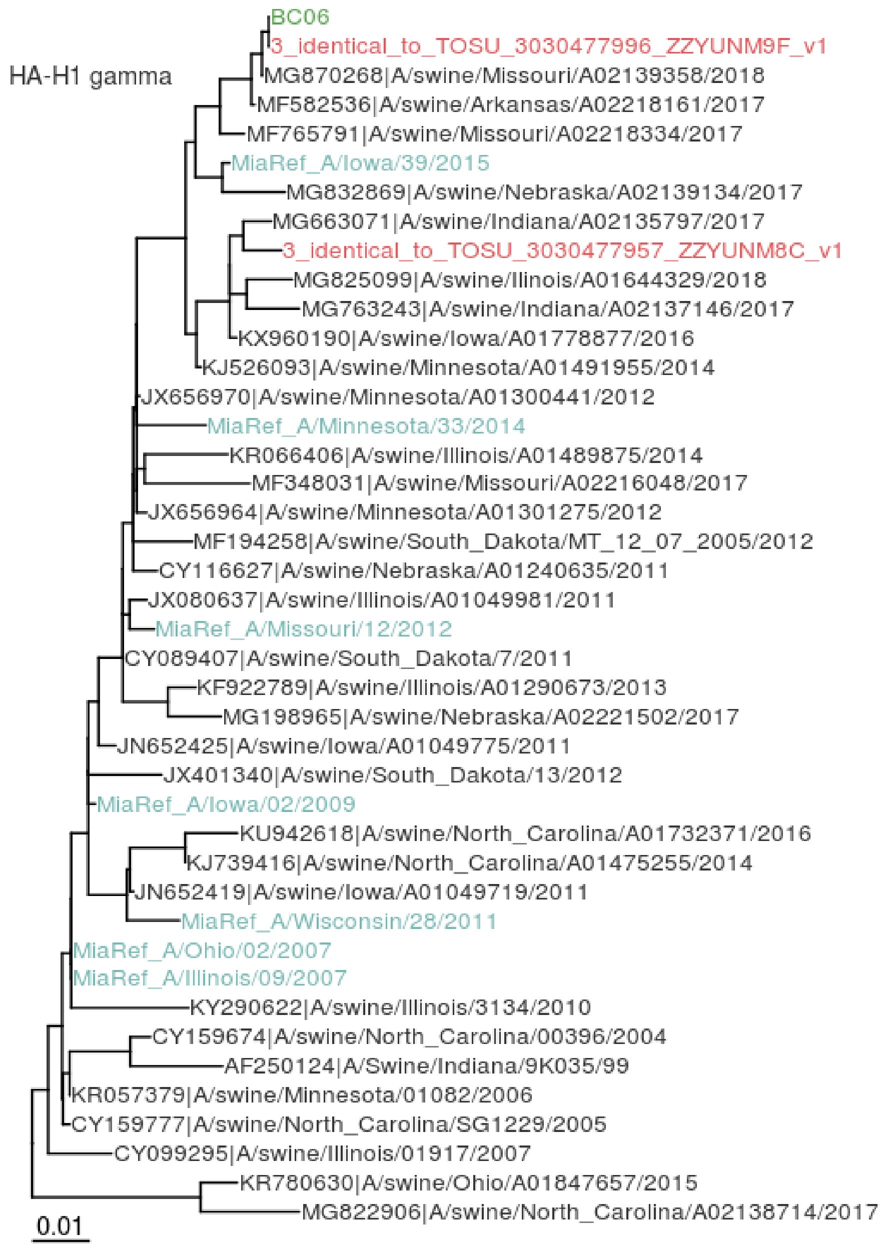


Post-field maximum-likelihood phylogenetic analysis: HA H1 gamma. In-field PCR amplicons sequenced by MiSeq are in green and additional swine IAV samples collected around the fair and laboratory-processed are in red. *Mia* references are in light blue. Diverse swine IAV sequenced from up-to-date USDA surveillance are in dark gray. Scale bar indicates average nucleotide substitutions per site.

**Figure S34**

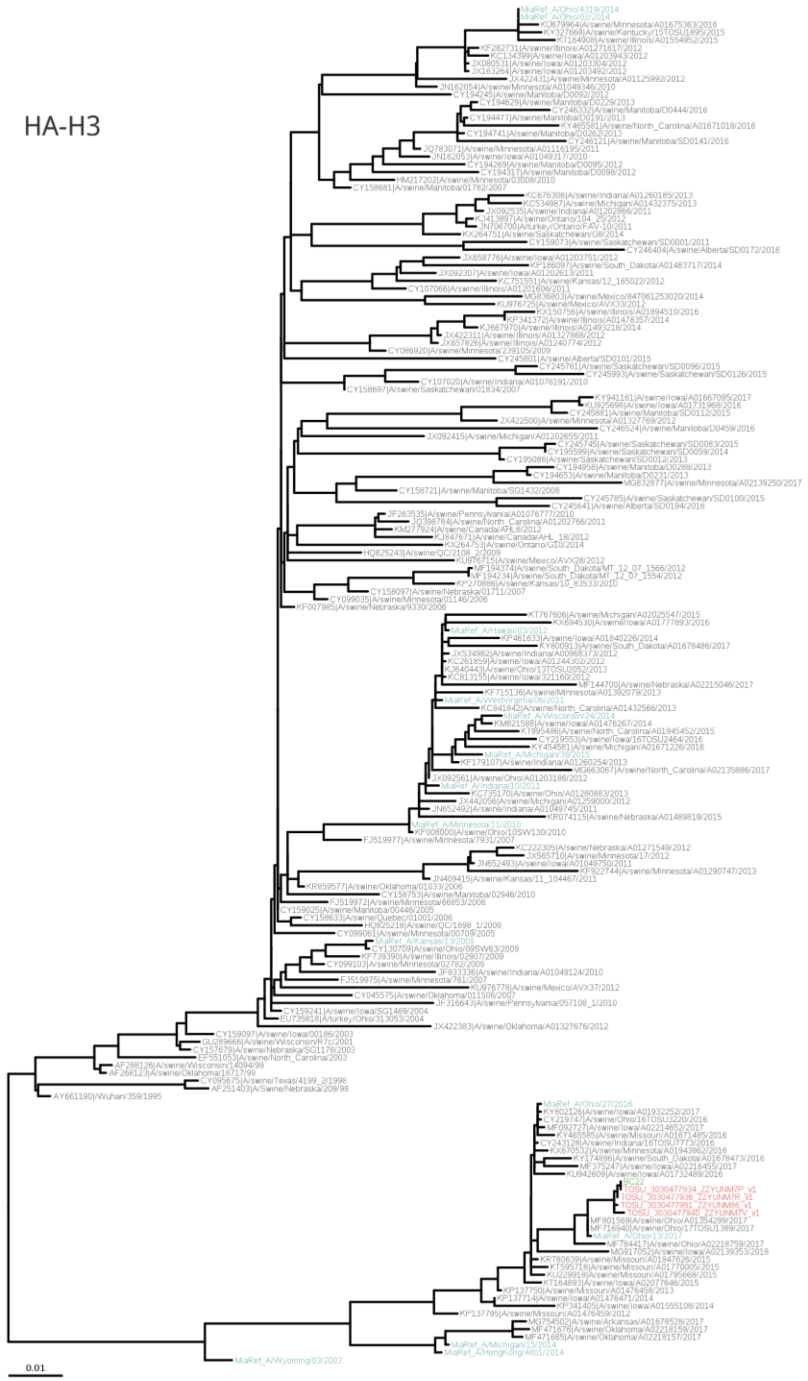


Post-field phylogenetic analysis: HA H3 2010 human-like. In-field PCR amplicons sequenced by MiSeq are in green and additional swine IAV samples collected around the fair and laboratory-processed are in red. *Mia* references are in light blue. Diverse swine IAV sequenced from up-to-date USDA surveillance are in dark gray. Scale bar indicates average nucleotide substitutions per site.

**Figure S35**

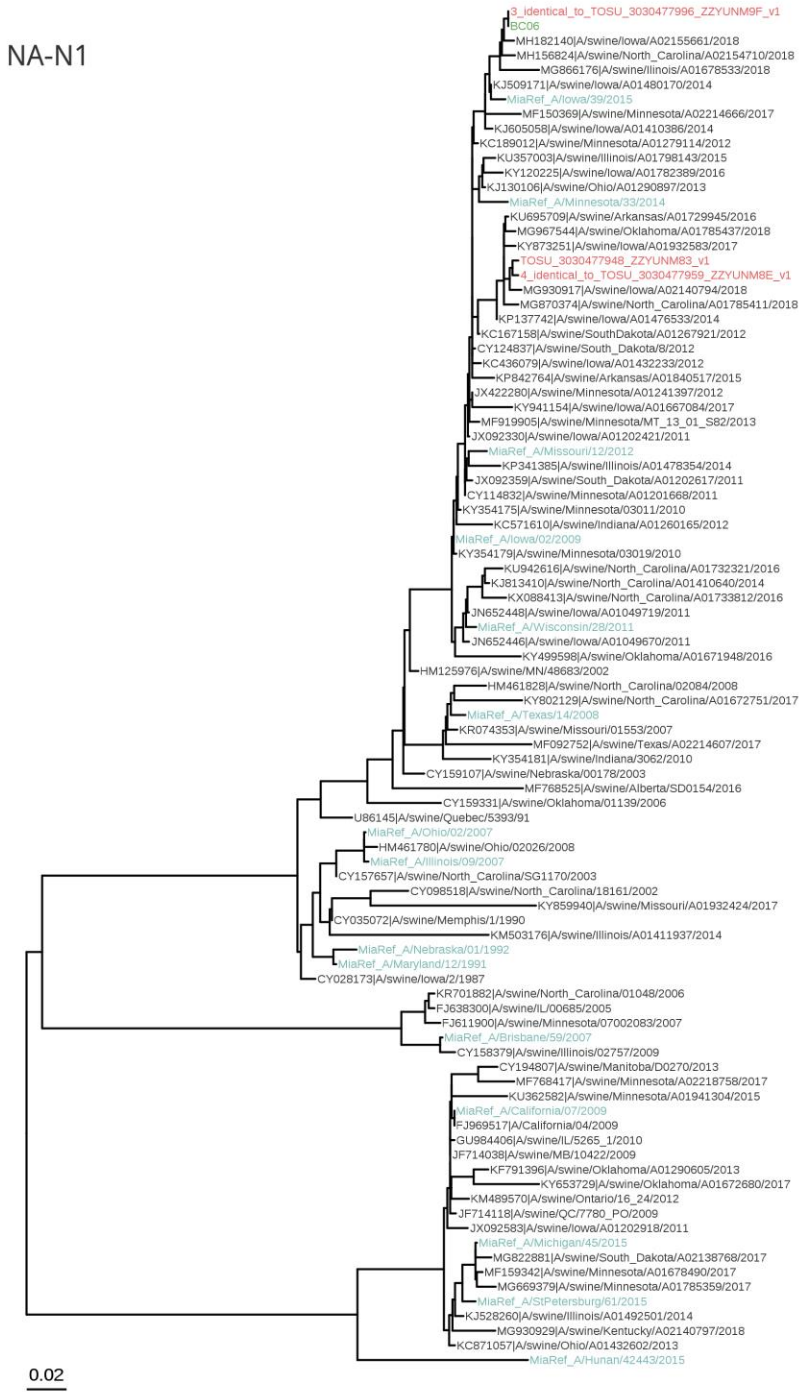


Post-field phylogenetic analysis: NA N1. In-field PCR amplicons sequenced by MiSeq are in green and additional swine IAV samples collected around the fair and laboratory-processed are in red. *Mia* references are in light blue. Diverse swine IAV sequenced from up-to-date USDA surveillance are in dark gray. Scale bar indicates average nucleotide substitutions per site.

**Figure S36**

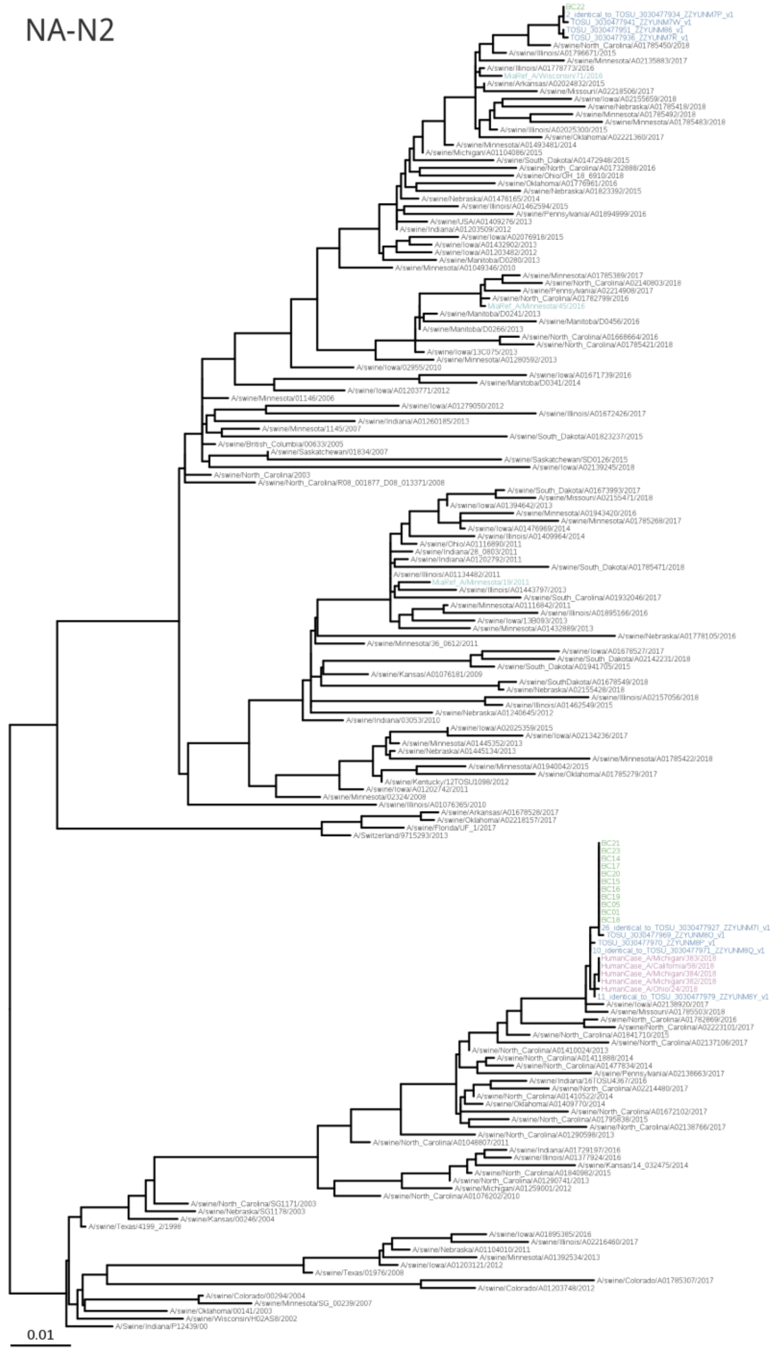


Post-field phylogenetic analysis: NA N2. In-field PCR amplicons sequenced by MiSeq are in green and additional swine IAV samples collected around the fair and laboratory-processed are in red. *Mia* references are in light blue. Diverse swine IAV sequenced from up-to-date USDA surveillance are in dark gray. Human cases of swine-variant viruses are in purple. Scale bar indicates average nucleotide substitutions per site.

**Figure S37**

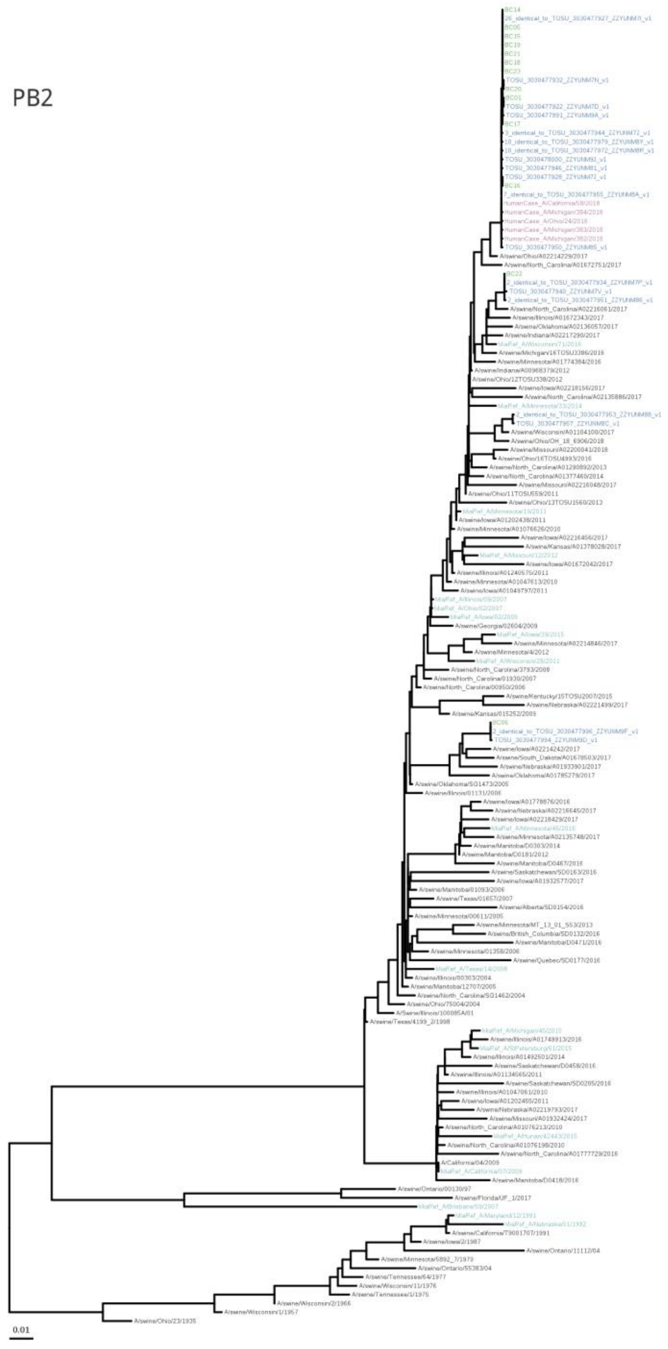


Post-field phylogenetic analysis: PB2. In-field PCR amplicons sequenced by MiSeq are in green and additional swine IAV samples collected around the fair and laboratory-processed are in red. *Mia* references are in light blue. Diverse swine IAV sequenced from up-to-date USDA surveillance are in dark gray. Human cases of swine-variant viruses are in purple. Scale bar indicates average nucleotide substitutions per site.

**Figure S38**

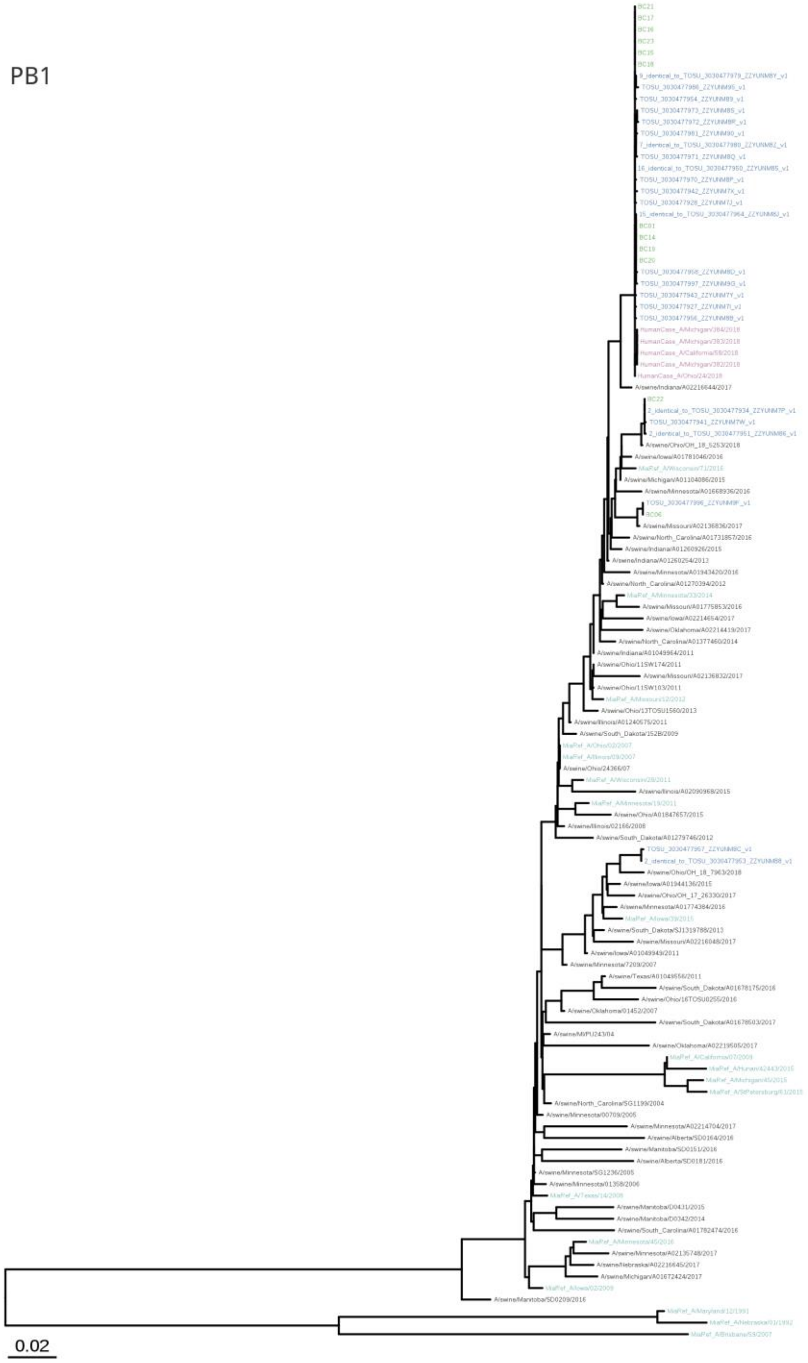


Post-field phylogenetic analysis: PB1. In-field PCR amplicons sequenced by MiSeq are in green and additional swine IAV samples collected around the fair and laboratory-processed are in red. *Mia* references are in light blue. Diverse swine IAV sequenced from up-to-date USDA surveillance are in dark gray. Human cases of swine-variant viruses are in purple. Scale bar indicates average nucleotide substitutions per site.

**Figure S39**

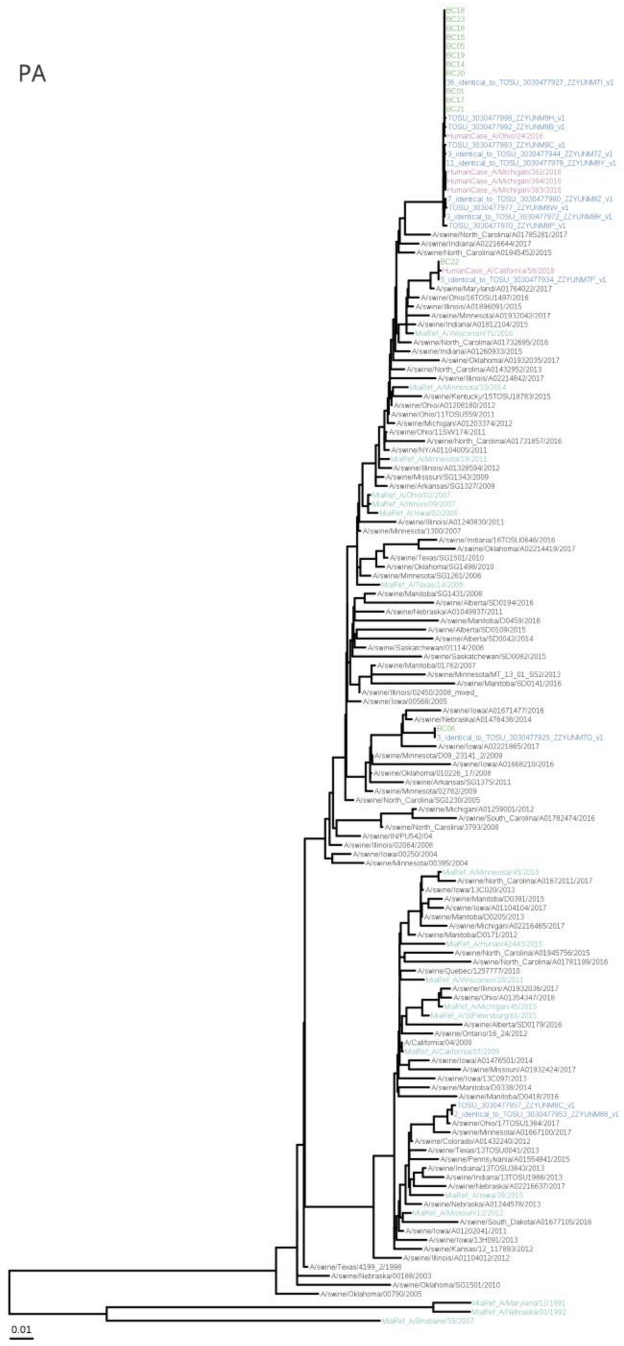


Post-field phylogenetic analysis: PA. In-field PCR amplicons sequenced by MiSeq are in green and additional swine IAV samples collected around the fair and laboratory-processed are in red. *Mia* references are in light blue. Diverse swine IAV sequenced from up-to-date USDA surveillance are in dark gray. Human cases of swine-variant viruses are in purple. Scale bar indicates average nucleotide substitutions per site.

**Figure S40**

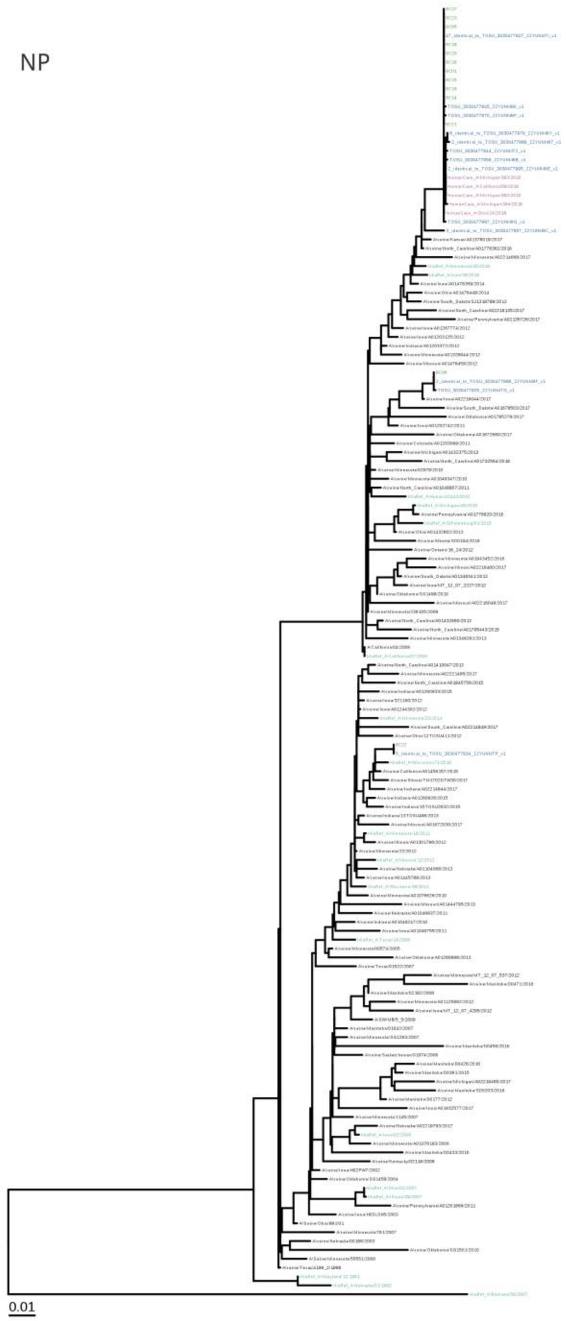


Post-field phylogenetic analysis: NP. In-field PCR amplicons sequenced by MiSeq are in green and additional swine IAV samples collected around the fair and laboratory-processed are in red. *Mia* references are in light blue. Diverse swine IAV sequenced from up-to-date USDA surveillance are in dark gray. Human cases of swine-variant viruses are in purple. Scale bar indicates average nucleotide substitutions per site.

**Figure S41**

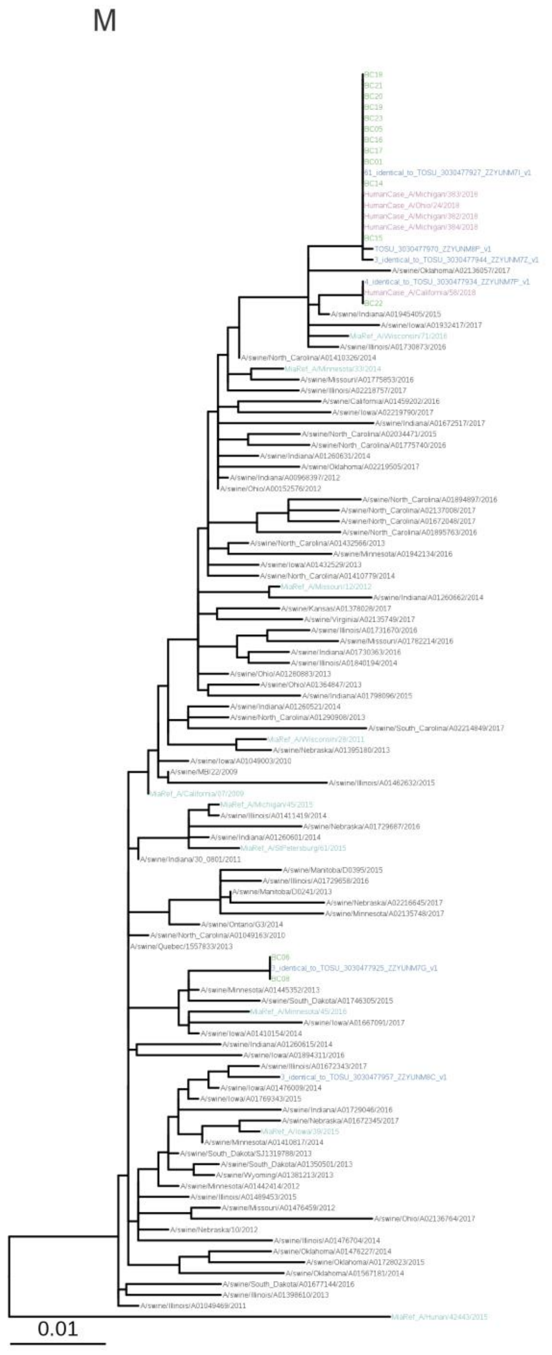


Post-field phylogenetic analysis: M. In-field PCR amplicons sequenced by MiSeq are in green and additional swine IAV samples collected around the fair and laboratory-processed are in red. *Mia* references are in light blue. Diverse swine IAV sequenced from up-to-date USDA surveillance are in dark gray. Human cases of swine-variant viruses are in purple. Scale bar indicates average nucleotide substitutions per site.

**Figure S42**

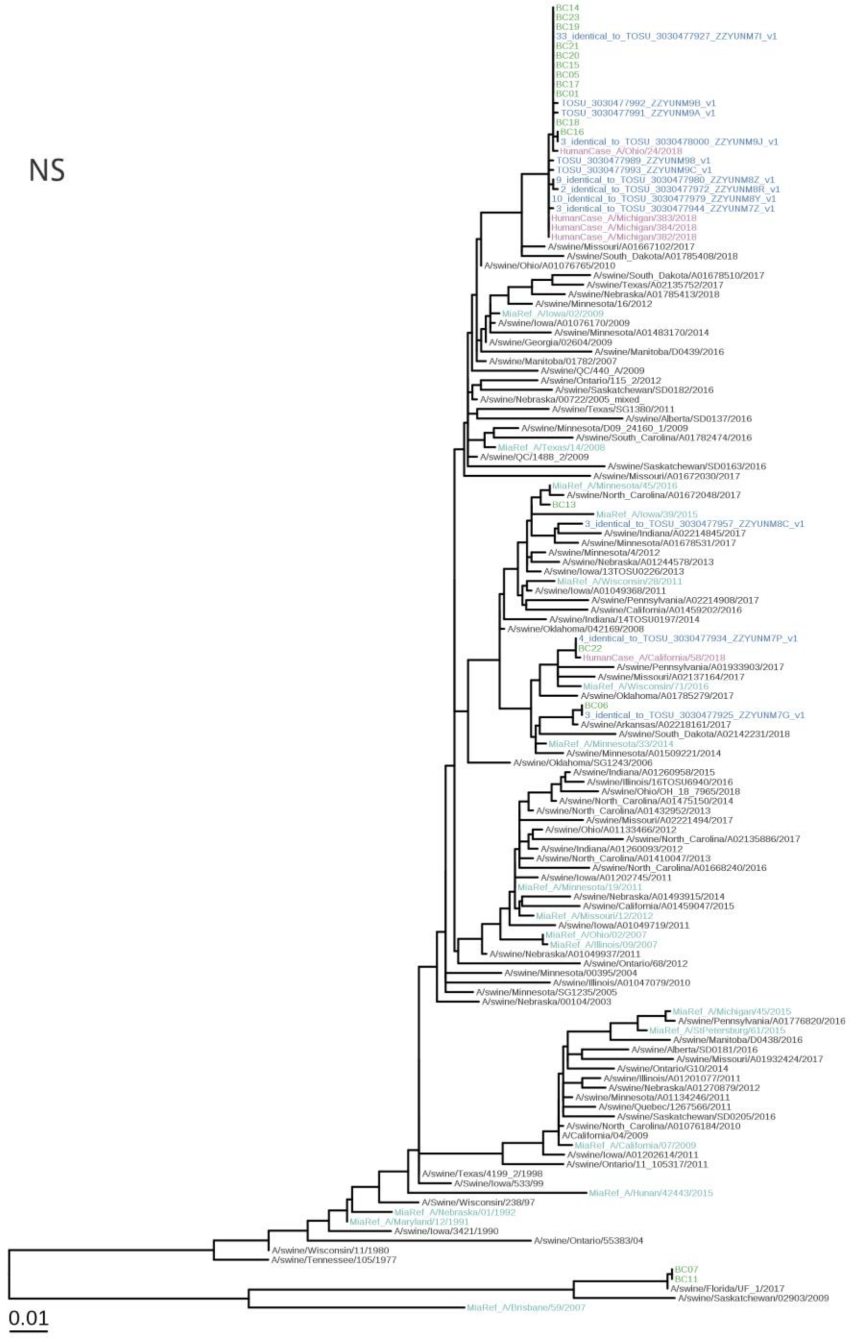


Post-field phylogenetic analysis: NS. In-field PCR amplicons sequenced by MiSeq are in green and additional swine IAV samples collected around the fair and laboratory-processed are in red. *Mia* references are in light blue. Diverse swine IAV sequenced from up-to-date USDA surveillance are in dark gray. Human cases of swine-variant viruses are in purple. Scale bar indicates average nucleotide substitutions per site.

**Figure S43**

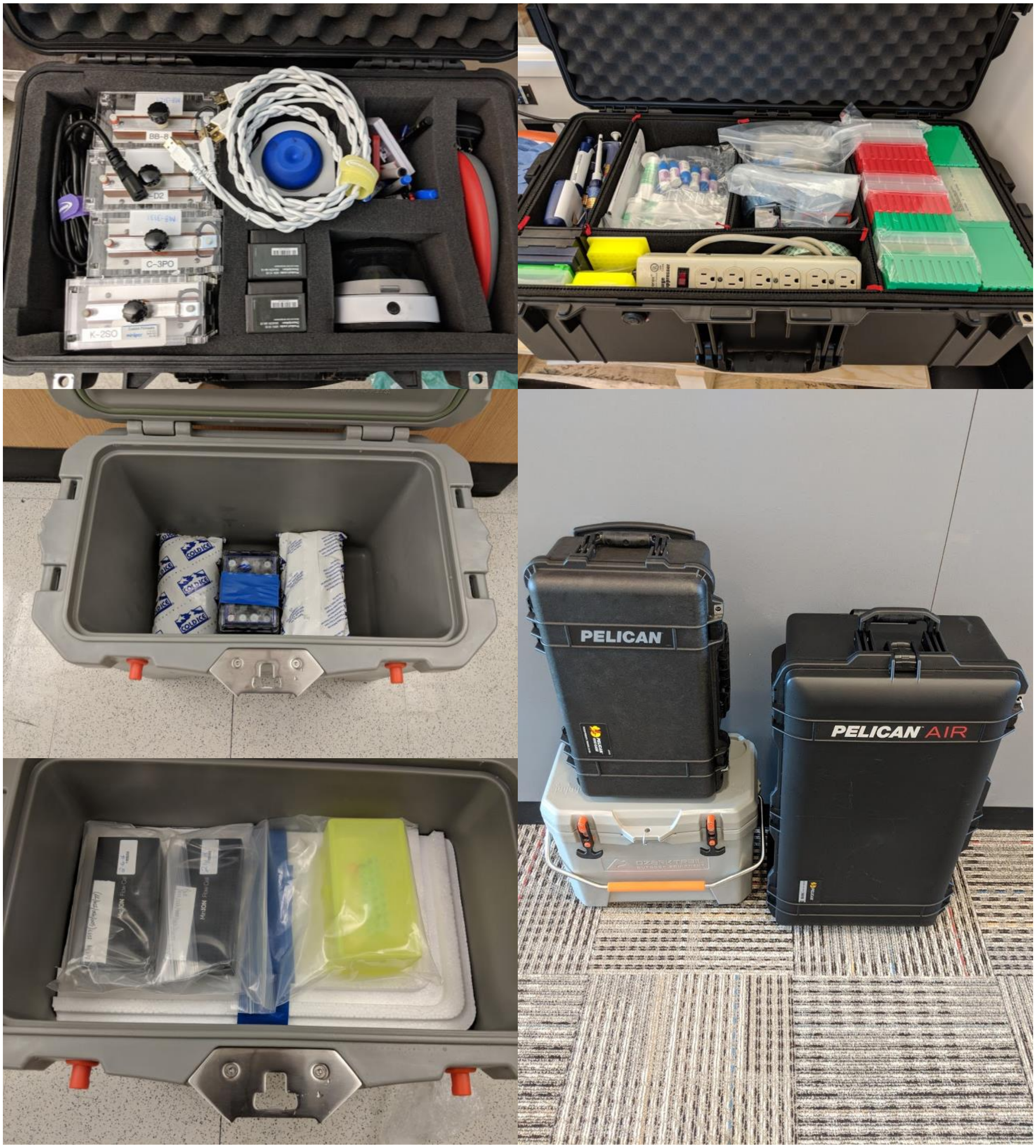


The assembled *Mia* pipeline occupies two suitcases and a cooler. The contents are listed in **Table S1**.

**Figure S44**

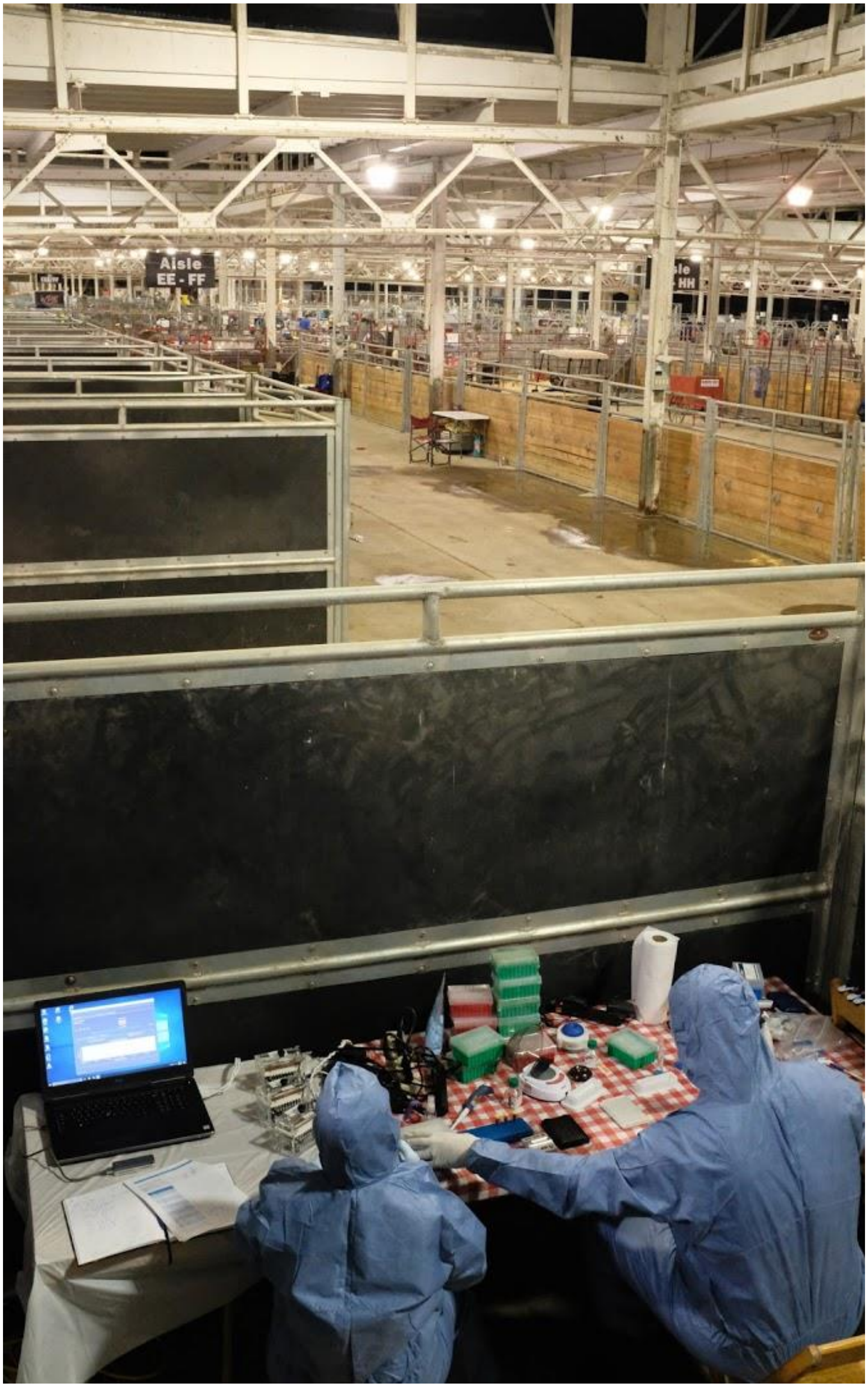


A photograph of the *Mia* setup mid workflow. We worked overnight in a vacant horse stall near the swine pins. From left to right: high performance laptop, minION, mini8 thermocyclers, Malania Wilson, power strips, tube racks and working space, pipette tips, centrifuge, vortex, paper towels, Matthew Keller, plastic consumables, pipettes hanging on a chair.
